# Cell-of-Origin, not Oncogenic Effect, Determines Desmoplastic Immune Exclusion in KRAS-Driven Liver Cancer

**DOI:** 10.64898/2026.03.24.711280

**Authors:** Chun-Shan Liu, Yu-Le Wu, Daria Komkova, Viktorie Gabrielova, Zeynep Cil, Trinh Kieu Dinh, Maximilian Zehender, Martin Schneider, Nugzar Lekiashvili, Philip Puchas, Darjus F. Tschaharganeh, Mathias Heikenwälder, Silvia Affo, Christoph Springfeld, Jan Pfeiffenberger, Conrad Rauber, Peter Sauer, Osman Öcal, Patrick Michl, Ângela Gonçalves, Felix J. Hartmann, Michael T. Dill

## Abstract

Intrahepatic cholangiocarcinoma (iCCA) and hepatocellular carcinoma (HCC) are the two most common primary liver cancers and share common risk factors. Yet they exhibit distinct oncogenic driver landscapes and fundamentally different tumor microenvironments (TME), with iCCA characterised by dense desmoplastic stroma that limits therapeutic efficacy. Whether these differences reflect oncogenic context or the developmental lineage of the cancer cell has remained unresolved.

Here, using syngeneic orthotopic murine models derived from CRISPR-engineered cholangiocyte and hepatocyte organoids each carrying *Trp53* deletion and *Kras^G12D^* mutation, we show that cell-of-origin, not oncogenic pathway activation, is the dominant determinant of TME architecture. Spatial proteomics of ∼390,000 cells reveals that cholangiocyte-derived tumors develop a stromal barrier of peripherally enriched αSMA^+^ cancer-associated fibroblasts (CAFs) that physically excludes immune cells and elevates PD-1/PD-L1 engagement, whereas hepatocyte derived tumors permit broader immune infiltration. Transcriptional variance partitioning confirms lineage as the primary source of gene expression divergence. Integrating murine and human transcriptomic and secretomic datasets, we identify LAMC2 and uPA as cholangiocyte lineage-specific secreted factors that trigger CAF activation. Genetic deletion of either factor markedly impairs iCCA formation *in vivo*. These findings establish that lineage-encoded secretory programmes create a desmoplastic and immune-excluded stroma and identify LAMC2 and uPA as functionally relevant modulators of TME in KRAS-driven iCCA.

## Introduction

Hepatocellular carcinoma (HCC) and intrahepatic cholangiocarcinoma (iCCA) are the two most common primary liver cancers and share common risk factors.[1,2] Although preclinical evidence demonstrates that hepatocytes can undergo biliary transdifferentiation under certain conditions,[3–6] both cancer types are defined by distinct lineage expression programmes associated with hepatocytes and cholangiocytes, respectively.

Beyond lineage identity, HCC and iCCA differ fundamentally in morphology, genetic landscape, and tumor microenvironment (TME).[7] A defining feature of iCCA is a prominent desmoplastic stromal reaction, characterised by extensive extracellular matrix (ECM) deposition driven by cancer-associated fibroblasts (CAFs).[8] iCCA also tends to show more aggressive growth behaviour and a greater propensity for extrahepatic metastasis than HCC. [9,10]

The desmoplastic stroma constitutes a major barrier to both chemotherapeutic efficacy and anti-tumor immunity.[11] Accordingly, iCCA is predominantly considered immunologically “cold”,[12–14] and responses to immune checkpoint inhibition, despite its recent inclusion in first-line systemic therapy regimens, remain limited.[15–17] Multiple CAF subtypes in iCCA have been shown to play tumour-promoting roles through diverse functional mechanisms,[18,19] and targeting stromal fibroblast recruitment and activation is under active investigation as a strategy to improve treatment efficacy.[11,20,21]

Why iCCA induces substantially more desmoplasia than HCC remains poorly understood.[8] One challenge is that the two cancer types differ not only in lineage but also in their oncogenic driver landscapes,[22,23] and oncogenic pathways are themselves known to shape the TME.[24,25] This makes it difficult to disentangle the contributions of lineage versus oncogenesis to the stromal phenotype. Compelling indirect evidence for a lineage-driven effect, however, comes from rare combined HCC-CCA (cHCC-CCA) tumors. Within a single monoclonal tumor sharing identical early driver mutations, desmoplastic stroma is maintained selectively in the CCA compartment, arguing against a purely oncogene-dependent mechanism.[26] However, existing preclinical iCCA models predominantly rely on hepatocyte transdifferentiation, which conflate cell-of-origin with oncogenic context and thereby prevent a systematic dissection of their independent contributions to cancer-stroma induction.

To address this directly, we leveraged organoid technology to engineer syngeneic, orthotopic murine models of iCCA and HCC in which lineage and oncogenic genotype are independently controlled. Using cholangiocyte-and hepatocyte-derived organoids carrying both *Trp53* deletion and *Kras^G12D^* mutation, we show that cell-of-origin is the dominant determinant of the stromal-rich, immune-excluded TME of iCCA. Through integrated transcriptomic and proteomic analysis across mouse and human datasets, we further identify LAMC2 and uPA as lineage-specific secreted factors that trigger CAF activation and are functionally required for iCCA formation *in vivo*.

## Results

### Cell-of-Origin Determines Tumor Identity in an Organoid-derived Liver Cancer Model of *Trp53* deletion and *Kras^G12D^* Mutation

To establish a syngeneic, orthotopic murine iCCA model (**Figure 1A**), we derived wild-type cholangiocyte organoids (chol-WT) from C57BL/6 mouse livers and used CRISPR-Cas9 to introduce *Trp53* deletion and *Kras^G12D^*mutation sequentially, generating chol-PK organoids. These two alterations are among the most commonly frequent somatic events in human iCCA (33.7% and 20.1%, respectively) and are known drivers of biliary malignant transformation.[27,28] Transformed organoids retained spherical morphology *in vitro* but, unlike their wild-type counterparts, proliferated in the absence of exogenous growth factor supplementation (**Figure 1B**). Both chol-WT and chol-PK organoids expressed high levels of biliary markers (*Krt7, Krt18, and Krt19*), with markedly lower expression of hepatocyte marker genes (*Afp, Alb, Aldob,* and *Hnf4a*) compared to hepatocyte organoids (**SFigure 1A**). Immunohistochemical (IHC) analysis confirmed a CK19^+^/HNF4α^-^ identity in both organoid lines, consistent with their cholangiocyte origin (**Figure 1C)**. CD44, a cancer stem cell marker, was upregulated in chol-PK but not chol-WT organoids (**Figure 1C**). Following orthotopic intrahepatic implantation into syngeneic C57BL/6 mice, chol-PK organoids gave rise to tumors with >90% (n=9/9, **SFigure 1B**). Histological analysis of explanted tumors with a diverse host-derived TME compartment (**Figure 1D**). IHC confirmed retention of CK19^+^/HNF4α^-^ pattern in the cancer cells (**Figure 1D**). Taken together, the histopathological features of our chol-PK tumors closely recapitulate key morphologic hallmarks of human iCCA.

**Figure 1.**
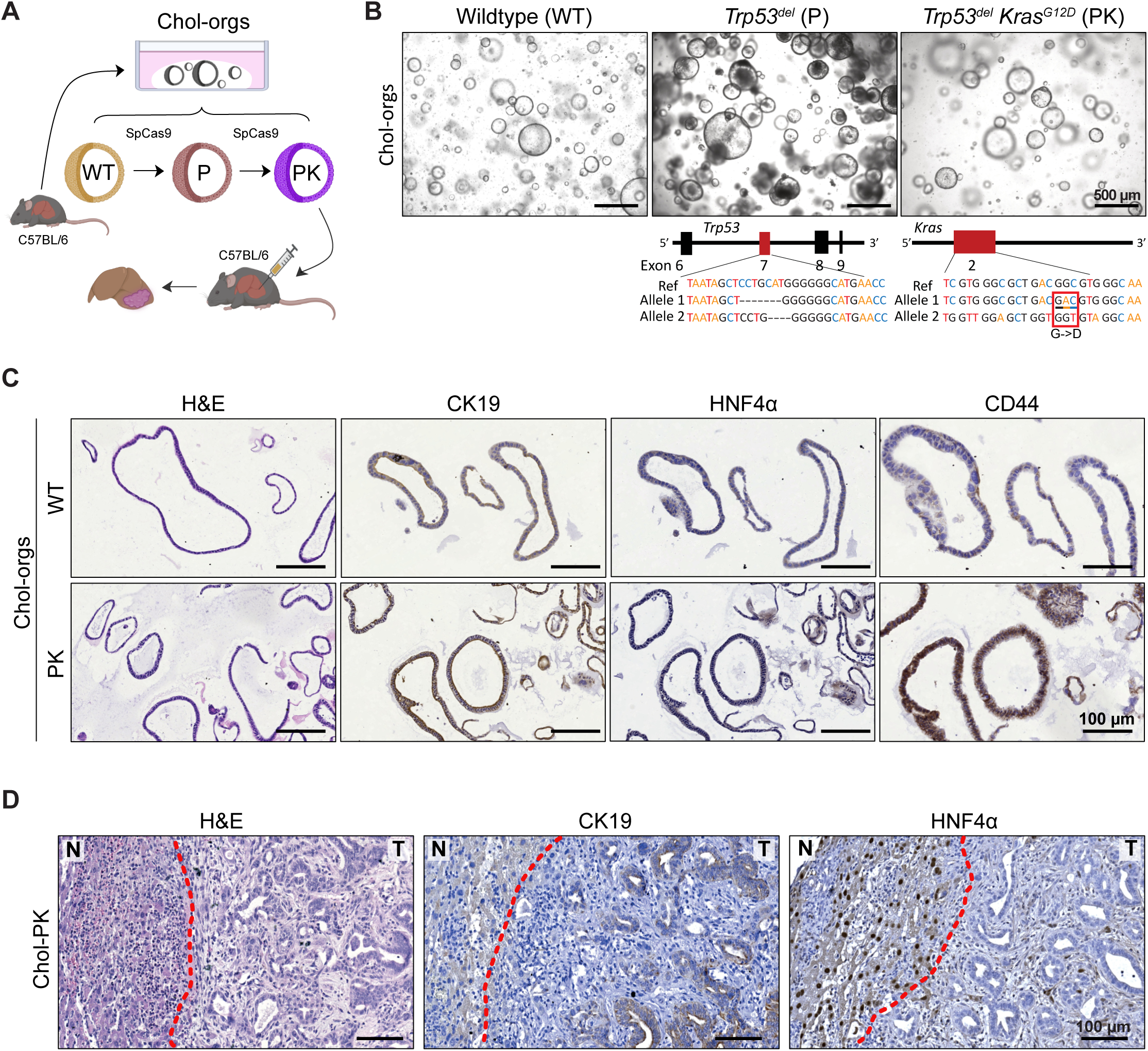
Cholangiocyte-derived organoids harboring *Trp53* deletion and *Kras^G12D^* mutation give rise to iCCA in a syngeneic orthotopic model. (A) Illustration of the syngeneic orthotopic CCA tumor model. Isolated wild-type chol-orgs were genetically engineered to harbor *Trp53* deletion and *Kras^G12D^*mutation by CRISPR/Cas9 and implanted intrahepatically. (B) Bright-field microscopic images of wildtype (WT) chol-orgs and with *Trp53* deletion (P) and *Kras^G12D^* mutation (PK). The CRISPR/Cas9-induced genetic modifications in *Trp53* and *Kras* genes are indicated below. (C) Representative stains of WT and PK chol-orgs indicating positive biliary lineage marker expression (CK19) and upregulation of CD44 in chol-PK orgs. (D) Representative histopathology images of liver tumors obtained upon orthotopic implantation of chol-PK resembling CCA. Stains as indicated. Red dotted line demarcates the boundary between non-tumor liver (N) and tumor (T). All scale bars in (B) indicate 500 μm, and in (C) and (D) 100 μm.

To independently control for lineage while maintaining the same oncogenic context, we next generated a hepatocyte-derived counterpart. Wild-type hepatocyte organoids (hep-WT) were derived from C57BL/6 mouse livers and engineered by lentiviral induction of *Trp53* deletion and *Kras^G12D^* overexpression to generate hep-PK organoids (**Figure 2A-B**). In contrast to the spherical morphology of chol-WT, organoids, hep-WT organoids formed dense, “grape-like” clusters, with correspondingly high expression of hepatocyte markers and low expression of biliary markers (**SFigure 1A**). IHC confirmed a CK19^-^/HNF4α^+^ identity in both hep-WT and hep-PK organoids, and CD44 upregulation was again restricted to the transformed line (**Figure 2C**). Following intrahepatic implantation, hep-PK organoids formed tumors with 75% penetrance (n = 9/12, **Figure 2D**). Macroscopic tumor size, mass and liver-to-body weight ratio did not differ significantly from chol-PK tumors (**SFigure 1B–E**). Despite comparable macroscopic appearance, histological analysis revealed strikingly distinct tumor morphologies. Hep-PK tumors displayed a trabecular growth pattern with densely packed cancer cells and minimal stromal area, reminiscent of human HCC and in contrast to the glandular, desmoplasia-rich architecture of chol-PK tumors. (**Figure 2E**). IHC confirmed retention of CK19^-^/HNF4α^+^ pattern, and none of the hep-PK tumors showed any histological features of iCCA (**Figure 2E**). These results establish that organoid lineage determines tumor morphology and differentiation, independent of shared oncogenic drivers.

**Figure 2.**
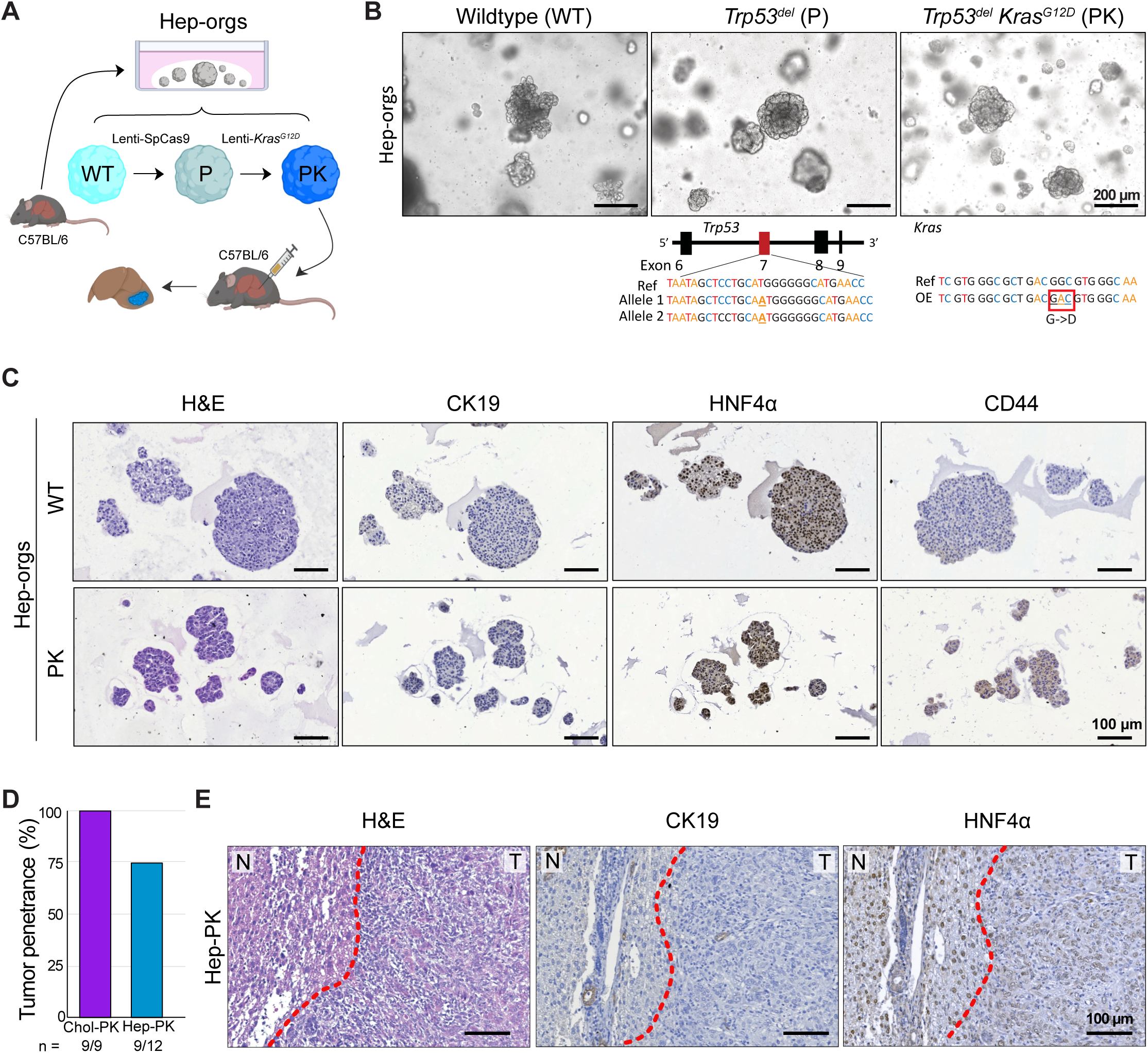
Hepatocyte-derived organoids harboring *Trp53* deletion and *Kras^G12D^* overexpression give rise to HCC in a syngeneic orthotopic model. (A) Illustration of the syngeneic orthotopic HCC tumor model. Isolated wild-type hep-orgs were genetically engineered to harbor *Trp53* deletion by CRISPR/Cas9 and *Kras^G12D^*via lentiviral transduction and implanted intrahepatically. (B) Bright-field microscopic images of wildtype (WT) hep-orgs and with *Trp53* deletion (P) and *Kras^G12D^*overexpression (PK). The CRISPR/Cas9-induced genetic modifications in the *Trp53* gene and the sequence of the overexpressing *Kras^G12D^* transgene are indicated below. (C) Representative stains of WT and PK hep-orgs indicating positive hepatocyte marker expression (HNF4α) and upregulation of CD44 in hep-PK orgs. (D) Bar plot of tumor penetrance of chol-PK and hep-PK organoids upon orthotopic tumor implantation. (E) Representative histopathology images of liver tumors obtained upon orthotopic implantation of hep-PK organoids resembling HCC. Stains as indicated. Red dotted line demarcates the boundary between non-tumor liver (N) and tumor (T). All scale bars in (B) indicate 500 μm, and in (C) and (E) 100 μm.

### Multiplexed Spatial Profiling Reveals Lineage-Dependent Immune and Stromal TME Architecture

To systematically characterise the spatial organisation of the TME, we performed multiplexed ion beam imaging (MIBI) with a 30-plex antibody panel targeting key lineage markers for immune cells, CAFs, endothelial cells and cancer cells (**Figure 3A**). Imaging focused on the tumor leading edge as the region of greatest immunological interest, encompassing ∼1.6 mm tissue zones extending from the adjacent liver into the tumor core (**Figure 3B**). A total of 408 fields of view (FOVs) were acquired from 4 chol-PK and 3 hep-PK tumors at 390 nm resolution, yielding a dataset of ∼390,000 cells with a mean of ∼46,000 cells per tumor. More than 75% of cells per section were confidently assigned to a defined cell type, and segmentation maps closely reflected H&E morphology, supporting the reliability of the classification (**Figure 3C-D, SFigure 2A–B**).

**Figure 3.**
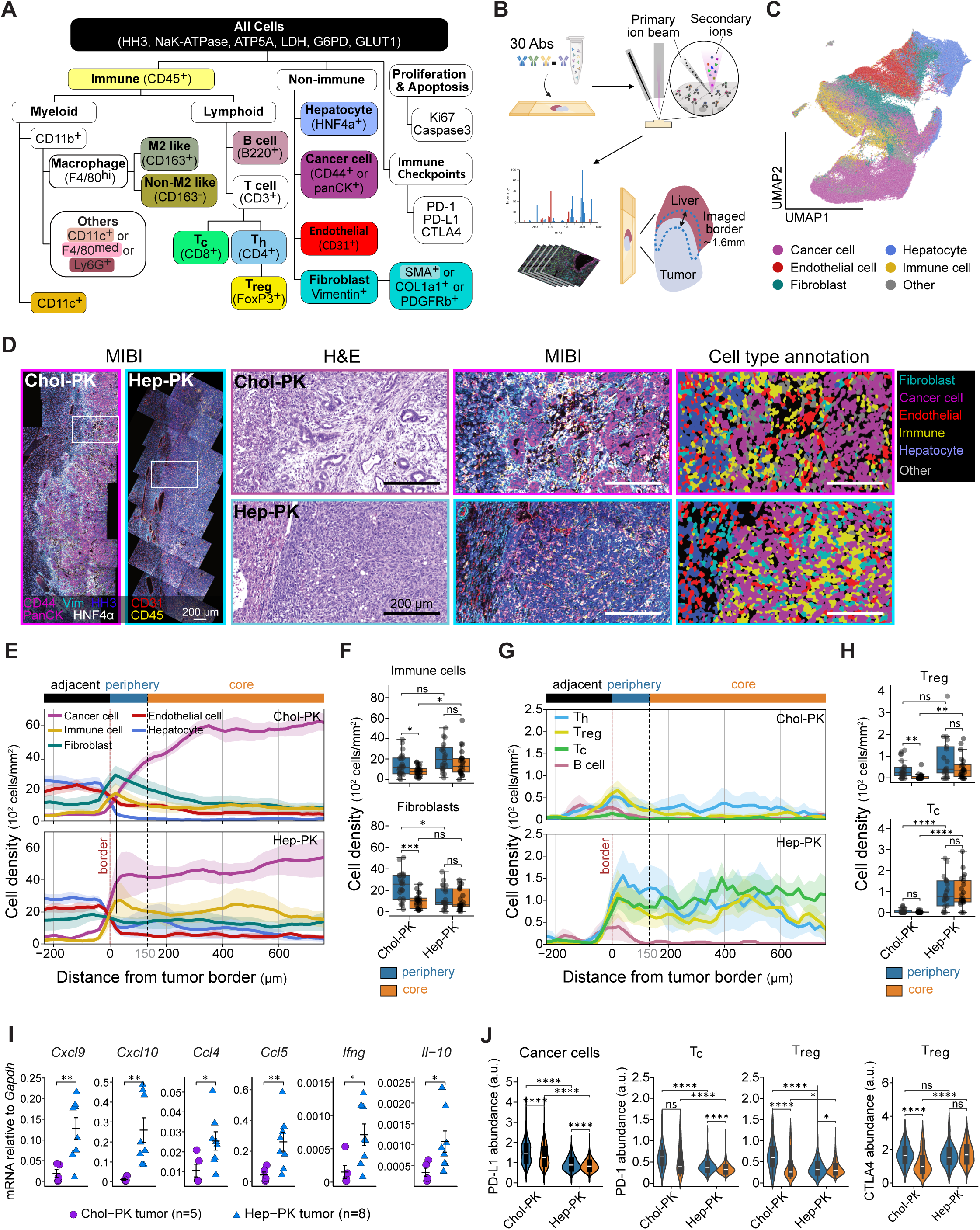
Multiplexed spatial profiling reveals a lineage-dependent, immune-excluded TME architecture in iCCA. (A) Hierarchical MIBI panel overview with 30 antibodies targeting cellular organelles, cell lineage and phenotypic markers, cell fate indicators, and immune checkpoint molecules. (B) MIBI workflow schematic and tumor region selection for imaging, spanning ∼1.6 mm from adjacent non-tumorous liver through tumor border into the tumor core. The tumor periphery was defined as the region extending 150 µm inward from the tumor boundary into the tumor. (C) UMAP projection of single cells based on lineage marker profiles. Each point represents one cell, comprising ∼390’000 cells from 4 Chol-PK and 3 Hep-PK tumors, colored by major cell lineages as indicated. (D) Representative H&E and MIBI images, and corresponding lineage annotations for chol-PK and hep-PK tumors. (E) Spatial distribution of indicated major cell types measured as cell density along an orthogonal axis of the tumor. (F) Box plot depicting immune cell and fibroblast densities separated by tumor periphery and core regions. (G) Spatial distribution of indicated lymphocyte subtypes measured as cell density along an orthogonal axis of the tumor border. (H) Box plot depicting cytotoxic T cells and Tregs separated by tumor periphery and core regions. (I) Messenger RNA expression of indicated pro-inflammatory cytokines and chemokines in chol-PK (n=5) and hep-PK (n=8) tumors. (J) Violin plots depicting spatial protein abundance of immune checkpoint molecules across single cells in the tumor periphery and core: PD-L1 in tumor cells, and PD-1 and CTLA-4 in Tc and Treg lymphocyte subtypes. Data from (E), (G) and (I) are presented as mean±SEM across tumor samples. In (F) and (H), each data point represents one tissue patch. Statistical significance was defined as * *p* ≤ 0.05, ** *p* ≤ 0.01, *** *p* ≤ 0.001, and **** *p* ≤ 0.0001.

Cell type distributions were quantified as mean density profiles along virtual axes perpendicular to the tumor border (**Figure 3E**). As expected, host hepatocyte density fell sharply at the tumour border while cancer cell density rose concordantly in both models, validating the spatial registration. Endothelial cell distributions were comparable between tumor types. By contrast, the distributions of CD45^+^ immune cells and vimentin-positive (Vim^+^) CAFs differed markedly (**Figure 3E–F**). In chol-PK tumors, CAF density peaked sharply at the border and declined towards the core, with immune cell density tracking a parallel but lower-amplitude profile. In hep-PK tumors, both CAFs and immune cells were more evenly distributed, with immune cells showing sustained infiltration into the tumor core. Quantitative comparison of a defined 150 µm peripheral zone versus the tumor core confirmed significant enrichment of immune cells, endothelial cells, and Vim^+^ CAFs in the periphery of chol-PK but not hep-PK tumors (**Figure 3F; SFigure 2C**). In summary, chol-PK tumors are characterised by peripheral accumulation of stromal and immune TME components with a paucity of immune infiltration in the core, whereas hep-PK tumors show broad, sustained immune infiltration throughout.

Consistent with this spatial segregation, chol-PK tumors showed a stromal-proliferative pattern at the periphery, marked by Ki67 positivity, with increasing apoptotic turnover towards the core, as indicated by cleaved Caspase-3 (Ccasp3) expression. Hep-PK tumors, by contrast, exhibited an immunoproliferative profile throughout (**SFigure 2D–E**).

To resolve immune cell subset differences, we next profiled the tumor immune microenvironment (TIME) in detail (**Figure 3G–H**). CD11c^+^ dendritic cells (DCs) and macrophages were present in both tumor types (**SFigure 2F–G**). In contrast, T cell subsets, including cytotoxic T (Tc), regulatory T (Treg), and T helper (Th) cells, mirrored the global TME distribution: accumulating at the border but largely excluded from the core in chol-PK tumors, while broadly infiltrating hep-PK tumors (**Figure 3G–H**). Despite comparable DC presence across models, T cell-attracting chemokines *Cxcl9*, *Cxcl10*, *Ccl4*, and *Ccl5* were expressed at significantly lower levels in chol-PK tumors (**Figure 3I**),[29] suggesting impaired T cell recruitment. *Ifng* and *Il10* mRNA were both elevated in hep-PK tumors, consistent with greater immune activation in this model (**Figure 3I**). At the protein level, chol-PK cancer cells exhibited significantly higher PD-L1 expression than hep-PK cells, and both Tc and Treg cells in chol-PK tumors displayed elevated PD-1, indicative of increased PD-1/PD-L1 axis engagement (**Figure 3J**).

Taken together, chol-PK tumors are characterised by a stromal-rich, immune-excluded microenvironment with elevated checkpoint engagement, whereas hep-PK tumors exhibit broad immune infiltration and greater immune activation. These differences are consistent with a dominant role of cell-of-origin in determining TME composition and immune architecture.

### Peripherally Enriched αSMA^+^ Myofibroblastic CAFs Form a Stromal Barrier at the iCCA Tumor Border

The peripheral accumulation of immune cells in chol-PK tumors prompted us to investigate whether a physical stromal barrier contributes to this immune-excluded architecture. Extracellular space, which was quantified as a surrogate for ECM content, excluding intraluminal areas, was significantly greater in chol-PK than hep-PK tumors (**Figure 4A**), corroborating H&E observations. CAFs are the primary source of ECM, providing structural support and influencing the TIME.[8,30,31] Supporting a broader relevance, CAF-associated genes (*ACTA2*, *COL1A1*, *FAP*, *S100A4*, and *VIM*) were significantly upregulated in human CCA relative to HCC in The Cancer Genome Atlas (TCGA) (**SFigure 3A**). To characterise CAF heterogeneity, we used MIBI expression of Vim, α-smooth muscle actin (αSMA), COL1A1, and PDGFRβ as a basis for iterative clustering, identifying six fibroblast subtypes (F1–F6) with distinct marker profiles (**Figure 4B, SFigure 3B**). αSMA^+^ CAFs were selectively enriched in chol-PK tumors, whereas hep-PK tumors were predominantly αSMA^-^ (**Figure 4C**). Spatially, αSMA^+^ CAFs accumulated at the tumor periphery, closely mirroring the immune cell distribution profile (**Figure 4D–E**), and showed elevated Ki-67 positivity in this region, indicating localised proliferative expansion (**Figure 4F**). αSMA is a marker for ECM-producing myofibroblastic CAFs (myCAFs), and consistent with this, approximately half of αSMA^+^ CAFs co-expressed COL1A1 (**SFigure 3B**, clusters F1–F2).[32]

**Figure 4.**
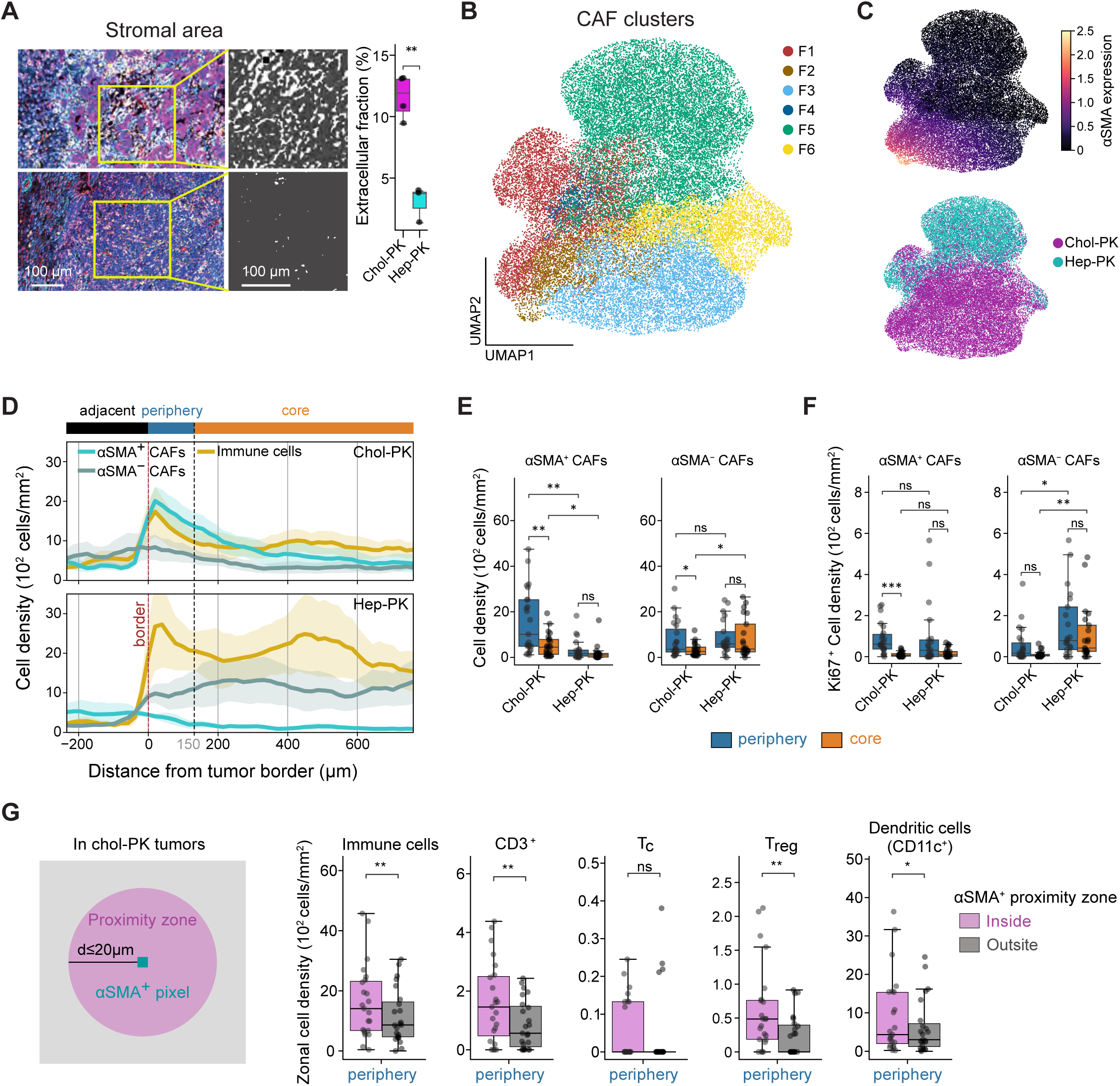
Peripherally enriched αSMA⁺ myCAFs form a stromal barrier associated with immune exclusion in iCCA. (A) Quantification of extracellular matrix area from multiplexed MIBI images. Left: representative images of chol-PK and hep-PK tumors. Middle: segmentation masks highlighting extracellular space, but excluding luminal areas. Right: Box plot of fraction of extracellular space. Each data point indicates one tumor. (B) UMAP projection of 38’152 annotated of single CAFs based on αSMA, Col1a1, and PDGFRβ protein levels. Each point represents one cell, comprising 38’152 CAFs from 4 chol-PK and 3 hep-PK tumors. Based on marker similarity, cells were classified into six clusters (F1–F6) based on αSMA, Col1a1, and PDGFRβ expression. (C) The same UMAP as in B indicating αSMA expression (top) and colored by tumor type (bottom), showing enrichment of αSMA⁺ CAFs in chol-PK tumors. (D) Spatial distribution of αSMA⁺ and αSMA⁻ CAFs alongside total immune cells measured as cell density along an orthogonal axis of the tumor border. (E) Quantification of αSMA⁺ and αSMA^-^ CAF densities in tumor periphery and core regions. (F) Proliferative activity of CAF subtypes, assessed by Ki-67⁺ cell density, showing peripherally enriched proliferative αSMA⁺ CAFs in chol-PK tumors. (G) Immune cell proximity to αSMA⁺ CAFs in chol-PK tumors. Left: schematic depicting a proximity zone defined as within 20 μm of αSMA⁺ pixels. Right: quantification of αSMA⁺ proximity zone for total immune cells, CD3⁺ T cells, Tc, Treg, and CD11c⁺ dendritic cells in the tumor periphery. Data from (D) is presented as mean±SEM across tumor samples. In (E), (F) and (G), each data point represents one tissue patch. Statistical significance was defined as * *p* ≤ 0.05, ** *p* ≤ 0.01, *** *p* ≤ 0.001, and **** *p* ≤ 0.0001.

To determine which immune cell types in chol-PK tumors are most spatially associated with αSMA^+^ CAFs, we measured the distance from each immune cell to the nearest αSMA^+^ pixel, defining proximity as ≤20 μm (∼1–2 cell diameters). In the tumor periphery, a significantly greater proportion of immune cells fell within this proximity zone than in the tumor core (**Figure 4G, SFigure 3C**). This spatial association was specific to T cells, particularly Tregs, and CD11c^+^ DCs, but was not observed for macrophages or other myeloid populations.

These findings support a model in which peripherally enriched αSMA^+^ myCAFs form a stromal barrier that restricts T cell entry, directly or indirectly, and thereby contribute to the immune-excluded architecture of chol-PK tumors.

### Lineage Remains Predominant Over Oncogenic Activation after Transformation

Because all non-cancerous TME components in our model are host-derived, the cancer organoids themselves must encode the lineage-specific signals that shape the distinct stromal architectures observed. HSCs are the principal source of CAFs in primary liver cancer,[18,21] and we therefore asked whether soluble factors secreted by chol-PK organoids could activate HSCs in a paracrine manner. Concentrated conditioned medium (CCM) from chol-PK organoids significantly induced expression of fibroblast activation markers *Acta2*, *Col1a1*, and *Fn1* in immortalised murine HSCs (mHSCs) compared to chol-WT CCM, whereas this difference could not be observed with CCM from hepatocyte-derived organoids (**Figure 5A**). These results implicate chol-PK-secreted factors as drivers of HSC activation and motivate their molecular identification.

**Figure 5.**
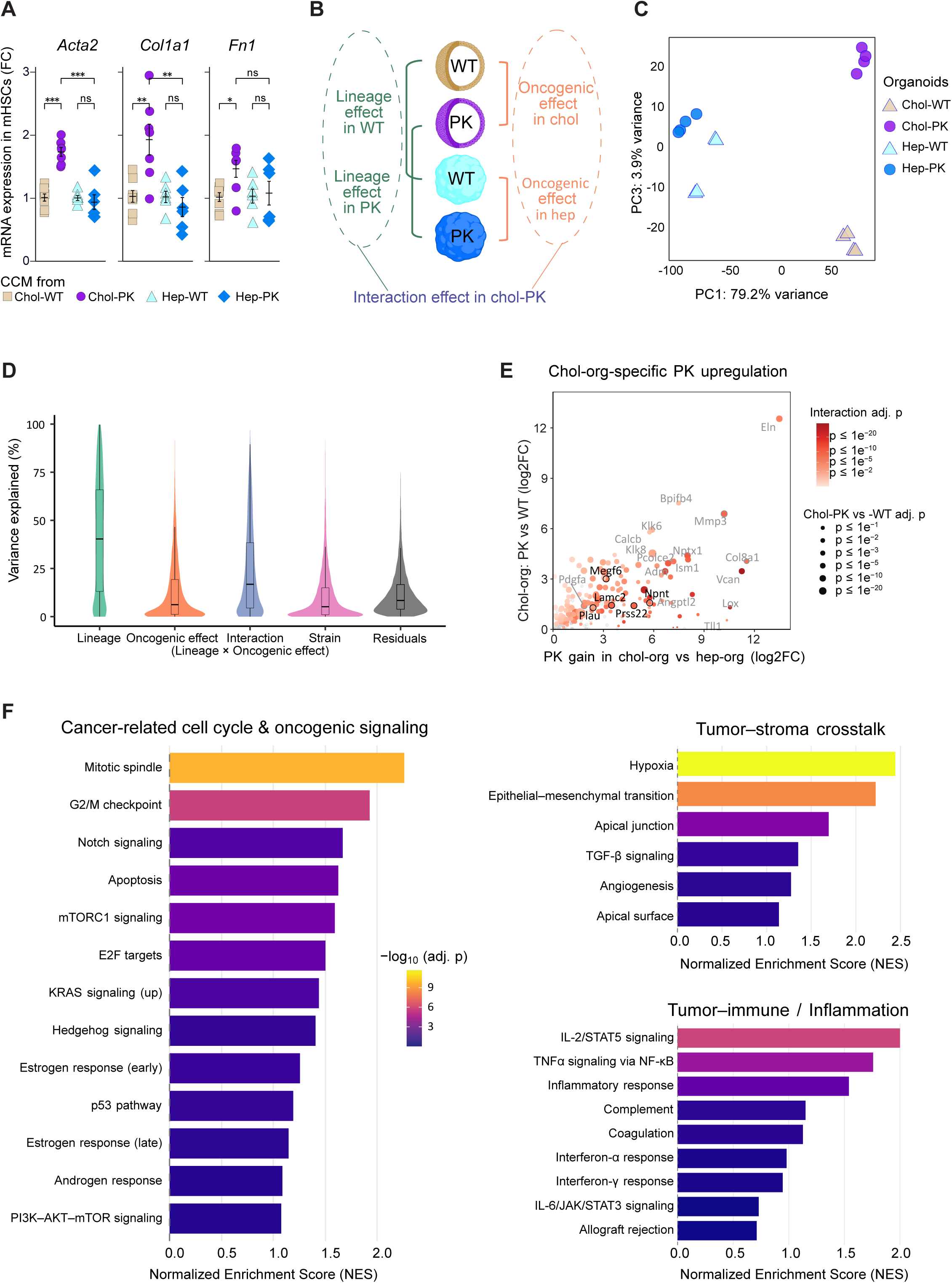
Cell-of-origin is the dominant determinant of transcriptional identity in *Trp53-deleted*, *Kras^G12D^*-mutant liver cancer organoids. (A) Messenger RNA expression of fibroblast activation markers in mHSCs treated for 24 hours with CCM from chol-or hep-derived organoids (WT or PK). (B) Schematic of multi-factorial transcriptome analysis in chol and hep organoids. Gene expression was modelled with lineage (chol vs hep), oncogenic effect (PK vs WT), and their interaction to identify lineage-specific PK effects. (C) Principal component analysis (PCA) plot of transcriptomes from indicated lines. Shown are PC1 (79.2% variance) versus PC3 (3.9%), which separate samples primarily by cell lineage and, to a lesser extent, oncogenic activation. (D) Distribution of variance in normalized expression of DEGs explained by lineage and oncogenic effect shown as violin plot with integrated box plot. For each gene, we fit a linear model on normalized counts with lineage, oncogenic effect, their interaction (lineage × oncogenic effect), and mouse strain as predictors. Residuals capture remaining variation. DEGs are the union across the four contrasts: chol-PK vs chol-WT, chol-PK vs hep-PK, chol-WT vs hep-WT, and hep-PK vs hep-WT. (E) Scatter plot showing upregulated DEGs in chol-PK organoids from multi-factorial analysis. Dot color indicates the significance of the interaction effect (adjusted p-value), and dot size reflects the significance of PK versus WT comparison in chol-orgs (adjusted p-value). (F) Gene Set Enrichment Analysis (GSEA) of Hallmark pathways specifically enriched in the lineage-dependent PK effect. The pathways selected for cancer-related cell cycle and oncogenic signaling, tumor-stroma crosstalk, and tumor-immune and inflammation with positive enrichment (i.e. PK effects stronger in chol-orgs) are shown.

To formally test the relative contributions of lineage and oncogenic activation to the transcriptional landscape, we performed bulk RNA-sequencing across all four organoid conditions (chol-WT, chol-PK, hep-WT, hep-PK; **Figure 5B**). PCA revealed that the dominant axis of variation (PC1) separated organoids by lineage, regardless of oncogenic status, i.e. *Trp53* deletion and *Kras^G12D^* mutation (**Figure 5C, SFigure 4A–B**). Strain differences among hepatocyte organoids accounted for PC2 (**SFigure 4A**), consistent with known inter-strain genomic variation,[33] while oncogenic status (WT vs. PK) contributed only to PC3, a substantially smaller source of variation than lineage. Variance partitioning confirmed these observations quantitatively: lineage accounted for the largest proportion of transcriptional variance, with oncogenic effects, and strain contributing substantially less (**Figure 5D, SFigure 4B–C**). Multi-factorial analysis for the interaction term “lineage × oncogenic effect” explained some variance, but these interaction effects largely reflected modulation of lineage-driven programs rather than independent oncogenic effects.

Accounting for these interaction effects, 3,235 genes were differentially expressed in chol-PK relative to chol-WT organoids (1,583 upregulated; **Figure 5E, STable 1**). Gene Set Enrichment Analysis (GSEA) revealed enrichment of cell cycle programmes (mitotic spindle, G2/M, E2F targets) and oncogenic signalling pathways (KRAS, PI3K–AKT–mTOR, NOTCH) consistent with malignant transformation (**Figure 5F, SFigure 4D**). Terms associated with tumor-stroma crosstalk, including EMT, TGF-β signalling, hypoxia, and angiogenesis, were also enriched, as were immune-related programmes including IL-2/STAT5 and TNFα–NF-κB signalling, which support T cell activation, and Interferon-γ response (**Figure 5F**). Conversely, enrichment of IL-6/JAK/STAT3 signalling, a driver of T cell exhaustion and immunosuppression, suggests that oncogenic activation concurrently induces immune evasion programmes.

Overall, these data reaffirm that in this controlled system, lineage is the dominant determinant of the transcriptional landscape, with oncogenic activation modulating, rather than overriding, pre-existing lineage-driven programmes.

### Multi-omics Integration Identifies LAMC2, uPA and BSSP-4 as CCA-specific Secreted Factors

Having established a paracrine role for chol-PK-secreted factors in mHSC activation, we next sought to identify the responsible molecules. We profiled the secretome and proteome of chol-WT and chol-PK organoids by mass spectrometry, using both concentrated conditioned medium (CCM) and cell lysate (CL) (**Figure 6A**). Integrated analysis of the murine transcriptome, secretome, and proteome, filtered against a reference list of human secreted proteins, identified 15 candidate factors enriched in chol-PK (**Figure 6B, SFigure 5A–B**). [33] To assess human relevance, we cross-referenced these candidates against proteome data from in-house human liver cancer organoid lines (**SFigure 5C**) and three published datasets (CCLE, TCGA, GSE26566) (**SFigure 5D–F**). [34–36] LAMC2, uPA (*PLAU*), and BSSP-4 (*PRSS22*) were consistently upregulated in CCA across all murine and human datasets and were selected for functional validation (**Figure 6C**).

**Figure 6.**
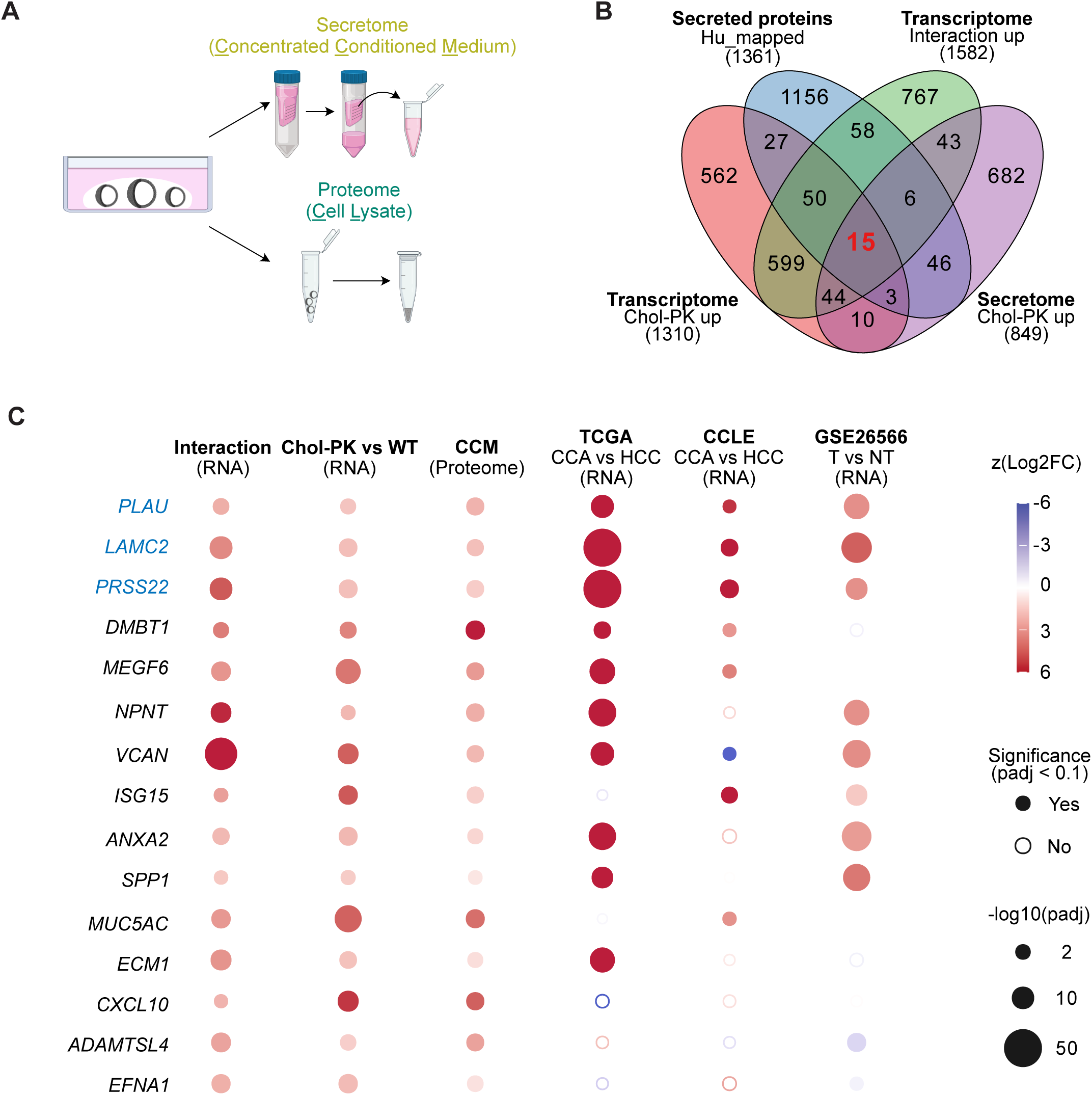
Multi-omics integration identifies LAMC2, uPA and BSSP-4 as CCA-specific secreted factors. (A) Schematic of the experimental workflow to collect material for secretome and proteome analysis. (B) Venn diagram showing the overlap of PK-induced transcriptional and secretory responses in chol-orgs. Intersections include PK-upregulated genes in chol-orgs from RNA-seq (chol-PK vs chol-WT, bottom-left), PK-upregulated proteins in chol-orgs from secretome (CCM, bottom-right), genes whose PK-induced expression is stronger in interaction effects (interaction contrast, upper-right), and proteins predicted to be secreted according to the Human Protein Atlas (upper-left). (C) Bubble plot showing cross-dataset support for candidate PK-associated genes identified by the integrative analysis in (B). Rows indicate 15 selected genes from (B), and columns indicate the indicated in-house murine result and public human datasets. Dot color encodes z-scored log2 FC (z(LFC). Filled circles indicate significant differences (padj < 0.1) and open circles indicate non-significant results. Dot size reflects statistical strength-log10(padj).

### LAMC2 and uPA affect HSC activation and impair iCCA formation *in vivo*

To test the individual contributions of LAMC2, uPA, and BSSP-4 to HSC activation, we pre-incubated chol-PK CCM with neutralising antibodies against each factor before applying it to mHSCs (**Figure 7A**). Anti-uPA and anti-LAMC2 antibodies each significantly reduced *Col1a1* expression, with anti-LAMC2 additionally suppressing *Acta2* (**Figure 7B**). Anti-BSSP-4 had no significant effect. Complementary gain-of-function experiments showed that recombinant uPA, but not BSSP-4, enhanced mHSC migration in a scratch assay (**Figure 7C–D, SFigure 6A**); full-length recombinant LAMC2 could not be tested owing to its large protein size.

**Figure 7.**
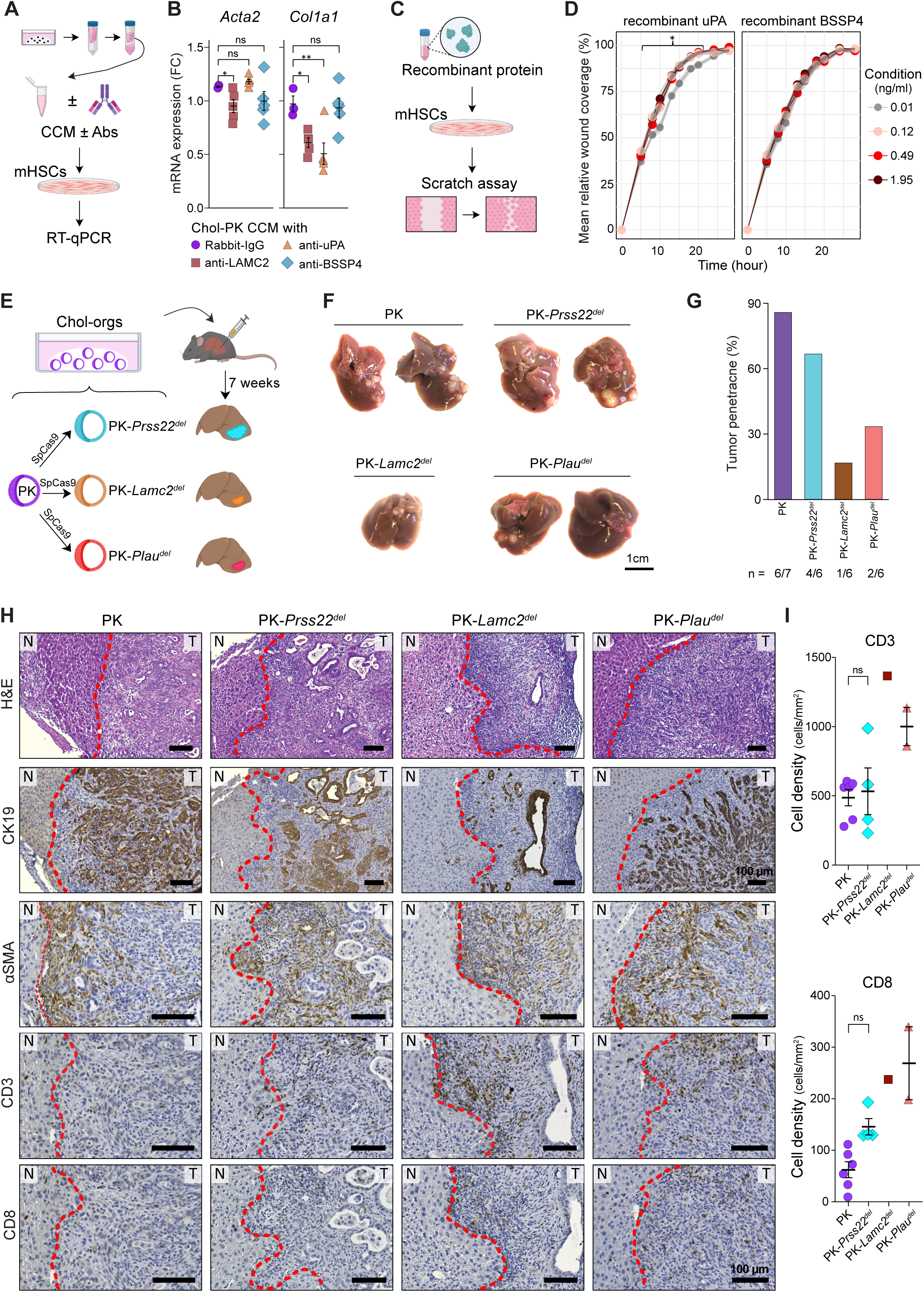
LAMC2 and uPA affect HSC activation and impair CCA formation *in vivo*. (A) Schematic overview of the *in vitro* fibroblast activation assay, where murine HSCs were treated with CCM from chol-PK and neutralizing antibodies. (B) Scatter plot indicating mRNA expression of *Acta2* and *Col1a1* in mHSCs treated with Chol-PK CCM and indicated neutralizing antibodies or control IgG. (n=5 from 2 independent experiments). (C) Schematic of scratch assay with mHSCs treated with low FBS (2%) media supplemented with respective recombinant protein. (D) Plot depicting mean relative wound coverage upon scratch injury over time with indicated recombinant protein at different dosage (n≥17 from 3 experiments). (E) Schematic of *in vivo* validation model: Chol-PK organoids were edited by CRISPR/Cas9 to generate *Prss22*-, *Lamc2*-, or *Plau*-deleted lines, which were orthotopically implanted into the liver and harvested at the same time point. (F) Macroscopic images of liver tumors 7 weeks after implantation of indicated lines. (G) Bar plot with respective tumor penetrance per line. (H) Representative histopathology images of liver tumors obtained upon orthotopic implantation of chol-PK and indicated KO lines. Stains as indicated. Red dotted line demarcates the boundary between non-tumor liver (N) and tumor (T). All scale bars are 100 μm. (I) Scatter plot depicting the total CD3^+^ and CD8^+^ T cell density per area per tumor for the indicated lines.

To test functional requirements *in vivo*, we generated chol-PK organoid lines with individual genetic deletions of *Lamc2*, *Plau*, and *Prss22* (**Figure 7E, SFigure 6B–C**). None of the knockout lines showed altered growth *in vitro*, indicating that these factors do not affect cell-intrinsic proliferative capacity (**SFigure 6D**). All three knockout lines and the chol-PK parental control were intrahepatically implanted under identical conditions and monitored for 7 weeks. At harvest, triggered when chol-PK had developed sizable tumors by IVIS imaging, tumor penetrance was markedly reduced in mice implanted with PK-*Plau^del^* (33%) and PK-*Lamc2^del^* (17%) organoids compared to parental chol-PK (86%) and PK-*Prss22^del^* (66%) (Figure 7G). The few tumors that did form in PK-*Plau^del^* and PK-*Lamc2^del^*mice were also substantially smaller than those in the other groups (**SFigure 6E**). Although formal statistical comparison was precluded by low tumor penetrance in some groups, PK-*Plau^del^* and PK-*Lamc2^del^*tumors showed a trend towards greater T cell and cytotoxic T cell infiltration, whereas PK-*Prss22^del^* tumors did not differ from parental controls (**Figure 7H–I**). In summary, LAMC2 and uPA, but not BSSP-4, activate HSCs *in vitro* and are functionally required for efficient iCCA formation *in vivo*.

## Discussion

The therapeutic consequences of oncogenic pathway activation are not universal. They are shaped by the cellular context in which they occur. Pan-cancer analyses have established that the dominant transcriptional programme of a tumor is primarily defined by cell-of-origin rather than oncogenic genotype and that oncogenic mechanisms observed in a certain cancer type cannot be fundamentally generalized.[34,37] Here, using cholangiocyte-and hepatocyte-derived organoid models carrying both *Trp53* and *Kras^G12D^* alterations, we demonstrate that lineage, not the shared oncogenic drivers, governs the morphology, stromal and immune architecture of liver cancer.

Our findings stand in contrast to some models in which *Kras* activation in hepatocytes induces iCCA.[28,38–41] Evidence suggests that biliary transdifferentiation of hepatocytes depends on the level of RAS or MYC signalling, [42,43] and that additional pathway activation, such as NOTCH or YAP1, or specific injury contexts promoting inflammation can further bias lineage outcome towards a biliary fate.[4,44] The heterogeneity across published models likely reflects the fact that existing systems vary lineage and oncogenic context simultaneously, making it difficult to distinguish cause from consequence. Our organoid-based system addresses this limitation by standardising the oncogenic context across both lineages, isolating cell-of-origin as the variable for tumor identity and TME formation.

In our system, hepatocyte-derived *Kras*-mutated organoids consistently gave rise to HCC upon implantation, without histological evidence of biliary differentiation in any tumour examined. This morphological fidelity was mirrored at the transcriptional level, where lineage-associated gene expression was maintained after oncogenic transformation. Whether enforced activation of biliary-fate determinants, such as NOTCH or YAP1, could override this lineage stability and induce iCCA in this system remains an open and worthwhile question.

Based on experience from preclinical models, the rarity of *KRAS*-mutated HCC relative to other gastrointestinal malignancies could be attributed to a lineage bias towards biliary cell transformation. Our findings raise a complementary hypothesis: that *Kras* activation in hepatocytes may confer a comparatively limited capacity for immune evasion relative to *Kras*-driven iCCA, which is characterised by a deeply immune-excluded microenvironment. If hepatocyte-derived *Kras*-mutant tumours are more immunologically vulnerable, this could contribute to their relative underrepresentation in clinical series. With KRAS inhibitors now entering clinical practice, a mechanistic understanding of how lineage context determines the immune consequences of KRAS activation will be essential for designing rational combination strategies with immunotherapy.[45,46] Our syngeneic organoid model provides a flexible platform to investigate these relationships.

The most striking TME differences between our two models were the depth of immune infiltration and the extent of ECM deposition, two features that were inversely correlated and lineage-dependent. Although total CAF numbers were comparable between iCCA and HCC tumors, MIBI analyses revealed a selective enrichment of αSMA^+^ myCAFs in iCCA. Spatially, these cells accumulated at the tumor periphery and closely apposed infiltrating lymphocytes, consistent with a physical stromal barrier that gates T cell entry into the tumor core. Supporting an active immunosuppressive role at this interface, peripheral Tregs exhibited higher PD-1 abundance than those in the tumor core. These features align with prior reports that αSMA^+^ CAFs are dominant stromal actors in iCCA and are linked to immune exclusion, desmoplastic remodelling, and poor outcome.[13,18,47,48] Further, αSMA^+^ CAFs can restrict CD8^+^ T cell access and confer resistance to checkpoint blockade in a breast cancer model,[49] rendering it an interesting cell population to target therapeutically.

Because all non-cancerous TME components in our model are host-derived, the cancer organoids themselves must be the source of signals that shape the distinct stromal architectures observed. Given that oncogenic genotype, implantation site, and host background are held constant across both models, lineage-encoded cancer cell-intrinsic programmes emerge as the most likely explanation for the differential CAF activation. Consistent with this, conditioned medium from chol-PK organoids was sufficient to activate HSCs, the principal CAF precursors in the liver.[18,21] Notably, canonical CAF-activating signals of the TGFβ family were not enriched in our multi-omics comparison of CCA-versus HCC-derived secretomes; instead, BSSP-4, uPA, and LAMC2 emerged as consistently CCA-enriched candidates.

Both uPA and LAMC2 have been identified as overexpressed in CCA,[36,50,51] and a recent proteogenomic characterization of CCA subtypes furthermore confirmed the relevance of our targets in the context of KRAS signaling, as both LAMC2 and uPA were found strongly enriched in the KRAS-mutated iCCA subtype.[52]

uPA and its receptor uPAR are central components of the plasminogen-activating pathway, upregulated across fibrotic diseases and multiple cancer types, with established roles in ECM remodelling, tumour invasion, and angiogenesis; inhibitors targeting this axis have entered clinical development.[53–55] LAMC2, a laminin chain component of the ECM, is overexpressed across multiple cancer types and associated with poor prognosis.[56,57] Beyond its structural role, the presence of EGF-like domains in LAMC2 confers cancer cell-autonomous signalling functions, including promotion of cell survival and epithelial-mesenchymal transition via EGFR and integrin pathways.[58] LAMC2 has been shown to drive progression in pancreatic adenocarcinoma[59,60] and to impair T cell migration in lung and oesophageal cancer models.[61] Notably, Zhang et al. recently reported that an intracellular full-length form of LAMC2 activates EGFR signalling in iCCA, an effect which has been attributed to the secreted form in prior work, and interpreted this as a cell-autonomous oncogenic function. [62] In our model, LAMC2 deletion did not affect organoid proliferation *in vivo*, but markedly impaired iCCA formation *in vivo*. This is consistent with a paracrine role, specifically, activation of HSCs and stromal remodeling. Whether the discrepancy with Zhang et al., is due to differences in oncogenic context (*Trp53*/KRAS versus AKT/YAP or AKT/NOTCH), cell-of-origin, or the possibility of an additional immune component not captured in that study, remains to be determined.

In conclusion, we establish that cell-of-origin, rather than shared oncogenic drivers, is the dominant determinant of desmoplastic immune exclusion in *Kras*-driven liver cancer. Using isogenic organoid-derived models in which lineage and oncogenic activation are independently controlled, we identify LAMC2 and uPA as lineage-encoded secreted mediators of CAF activation that are functionally required for iCCA formation. These findings argue that oncogenic pathway activity must be interpreted within the lineage context of the cancer cell, a consideration of direct relevance to cancer-agnostic therapeutic strategies targeting KRAS. Future work dissecting the interplay between cell-of-origin, oncogenic activation, and the tumorigenic niche will be essential to identify cancer subtype-specific vulnerabilities and exploit them therapeutically.

## Acknowledgements

The authors are very grateful to all patients and their families, who provided biomaterial for this research project. The authors thank Robert Vanner, Sven Truxa, Kassandra Hoetzel, and Carolin Brombach for the initial MIBI mouse panel design, and thank Mei-Ju May Chen, Li-Chin Wang and Pei-Chi Wei for support with RNAseq data interpretation. The authors thank Jutta Mohr, Petra Klöters-Plachky, Farzaneh Kashfi, Sarah Böhn, and Sabrina Schweiggert for outstanding technical support, and want to acknowledge the excellent services provided by the Center for Preclinical Research, Light Microscopy, Proteomics and Next Generation Sequencing Core Facilities of the DKFZ.

## Author Contributions

Conceptualization: C.-S.L., M.T.D.; Methodology: C.-S.L., Y.-L.W., D.K.T., N.L., P.P. M.T.D.; Investigation: C.-S.L., Y.-L.W., Z.C., V.G.; Validation: C.-S.L., Y.-L.W., D.K., V.G., M.T.D.; Formal Analysis: C.-S.L., Y.-L.W., D.K., M.Z., V.G., M.S., M.T.D.; Software: Y.-L.W., D.K., M.Z., M.S.; Resources: M.H., S.A., C.S., J.P., C.R., P.S., O.Ö., P.M., F.J.H., M.T.D.; Writing – Original Draft: C.-S.L., M.T.D.; Writing – Review & Editing: all authors; Visualization: C.-S.L., Y.-L.W., D.K., M.Z.; Supervision: A.G., F.J.H., M.T.D.; Project Administration: M.T.D.; Funding Acquisition: M.T.D.

## Materials and Methods

### Composition of organoid media

L-WRN conditioned media were generated as described previously.[63] Basal media was based on advanced DMEM/F12 (ThermoFisher Scientific, #12634010) supplemented with 1% (vol/vol) penicillin/streptomycin (ThermoFisher Scientific, #15140122), 1% (vol/vol) Glutamax (ThermoFisher Scientific, #35050038), 20 mM HEPES (ThermoFisher Scientific, #35050038), 1:25 B27 supplement (without vitamin A) (ThermoFisher Scientific, #12587010), 1:50 N2 supplement (ThermoFisher Scientific, #17502048), and 20 mM Nicotinamide (Sigma-Aldrich, #N0636). Organoid media was a 1:1 mixture of L-WRN conditioned media and basal media. Cholangiocyte minus growth factor media (chol GF-media) was based on organoid media supplemented with 100 µg/ml Primocin (Invivogen, #ant-pm1). Cholangiocyte plus GF media (chol GF+) media was based on chol GF-media plus 50 ng/ml murine EGF (ThermoFisher Scientific, #31509-1mg) and 50 ng/ml human HGF (ThermoFisher Scientific, #10039-500). Hepatocyte minus GF (hep GF-) media was based on organoid media supplemented with 1.25 mM N-acetyl-L-cysteine (Sigma-Aldrich, # A9165), 10 nM Gastrin (Tocris, #3006), 3 µM CHIR99021 (Tocris, #4423), 1 µM A83-01 (Tocris, #2939-10mg), 100ng/ml human TNF-a (R&D Systems, #7754BH-005), and 100 µg/ml Primocin. Hepatocyte plus GF (hep GF+) media was based on hep GF-media plus 50 ng/ml murine EGF, 25 ng/ml human HGF, 50 ng/ml FGF7 (ThermoFisher Scientific, #10019-250), and 50 ng/ml human FGF10 (ThermoFisher Scientific, #10026-500). Human liver organoid media was based on organoid media supplemented with 1 mM N-acetyl-L-cysteine, 10 nM Gastrin, 5 µM A83-01, 10 µM Forskolin (Tocris, Cat#1099-10mg), 25ng/ml human HGF, 50ng/ml human EGF, 100 ng/ml human FGF10 and 100 µg/ml Primocin. 10 µM Y27632 (Holzel, #TMO-T1725) was added in organoid media for passage.

### Murine liver organoid isolation and culture

Wild-type cholangiocyte and hepatocyte organoids were isolated from 4-6 week-old C57BL/6J (cholangiocyte and hepatocyte organoids) and C57BL/6N (hepatocyte organoids) mice, adapting a published protocol.[64] In short, after euthanasia via cervical dislocation and midline laparotomy, mouse livers were immediately perfused via portal vein by a pre-warmed (37°C) perfusion buffer (50 mM EDTA, 10 mM HEPES in 1X HBSS [GIBCO]) at 6 mL/min for 2 min, then 8 mL/min for another 2 min, followed by Collagenase IV digestion buffer (Collagenase IV:150 U/mL Invitrogen cat# 17104019; in 1X HBSS with 1.25 mM CaCl₂, 4 mM MgCl₂, 10 mM HEPES) for 5–10 min at 8 mL/min. The digested liver was then excised. After removal of the gallbladder, the liver capsule was removed, and the tissue gently agitated to release hepatocytes. The hepatocytes were then collected and washed by centrifugation at 30×g for 5 min 3 times in ice-cold resuspension buffer (2% FBS (ThermoFisher Scientific, #10270106), 1.25 mM CaCl₂, 4 mM MgCl₂, 10 mM HEPES, 5 mM glucose in 1X HBSS).

To obtain cholangiocytes, the remaining liver scaffold underwent sequential digestions with collagenase IV buffer (150 U/mL), accutase, and 0.25% trypsin for 30, 30, and 20 min respectively at 37°C. Cells were collected after each step, filtered (100 μm), washed, and resuspended in ice-cold resuspension buffer. The suspensions were then combined and spun at 50 G for 5 min at 4°C. The supernatant was then transferred and spun at 300 G for 5 min at 4°C. The cell pellet was then resuspended in basement membrane extract (BME, Cultrex Reduced Growth Factor Basement Membrane Extract, Type 2, R&D Systems, #3533-010-02) and plated in 24-well plates (50 μl droplets per well). After solidification, cholangiocyte BME droplets were overlaid with 500 μl of mouse chol GF+ media. After 7-14 days, outgrowing cholangiocyte organoid clones were picked for further passage.

For hepatocytes, the cell pellet was resuspended in 4.8 mL of 40% iodixanol (Optiprep) diluted in resuspension buffer (1:2.2 cell:iodixanol ratio). The suspension was overlaid with 3 mL of 18% iodixanol and 0.5 mL resuspension buffer, then centrifuged at 500×g for 25 min at 4°C with reduced deceleration. The top cell layer, enriched for viable hepatocytes, was collected, mixed with wash medium (DMEM (ThermoFisher Scientific, #31966021) with 1% FBS), and washed at 30×g for 5 min. The cell pellet was resuspended in BME (~30,000 cells per 50 µl BME) and plated in 24-well plates (50 μl droplet per well). Hepatocyte BME droplets were overlaid with 500 μl of hep GF+ media.

For further passaging, organoids were mechanically disrupted by 1ml syringe with 25G needle and cultured in the indicated media. Media was changed two to three times per week.

### Human Liver Cancer Organoid Generation and Culture

For the isolation of liver cancer cells, spare tissue from either liver tumor biopsy specimens or material from endoscopic retrograde cholangiography were obtained during routine diagnostic work-up at the Heidelberg University Hospital. Written informed consent was obtained from all patients, and the study was approved by the ethics committee of Heidelberg University (ethics permit numbers S-255/2021 and S-751/2020). The liver tumor biopsy pieces were cut into small pieces with a scalpel and incubated with a digestion solution (EBSS containing CaCl_2_ and MgCl_2_ supplemented with collagenase D [2.5 mg/ml] and DNase I [0.1 mg/ml]) for approximately 15 minutes at 37°C in a cell incubator. The digestion was then stopped by adding ice-cold DMEM with 1% FBS. The digested pieces were filtered through a 70 µm nylon cell strainer and centrifuged with 300 G for 5 minutes at 4°C. Both the digested pellet and remaining pieces in the filter were separately cultured in BME at 50 µl droplets. After hardening of the BME dome, human liver organoid media was added. For further passaging, organoids were mechanically disrupted by 1ml syringe with 25G needle and cultured in the indicated media. Where necessary, outgrowing tumor organoids were separated from wildtype cholangiocyte organoids by picking with a pipette. Media was changed two times per week.

### Cloning of sgRNA and lentiviral vector constructs

To create sgRNA constructs, pX458(BB)-GFP and pX459(BB)-puro backbone plasmids were used (Addgene, #48138 and #62988).[65] The pX458-sgRNA(*Trp53*) plasmid was generated by integrating established *Trp53* sgRNA oligos [66] into the pX458 vector. sgRNA oligos for *Plau*, *Lamc2* and *Prss22* were designed using Benchling (https://benchling.com) and cloned into pX459(BB)-puro. The sgRNA(Kras)-KrasG12D_HDR plasmid was generated by modifying the parental plasmid AAV-KPL (Addgene #60224).[66] Briefly, AAV-KPL was digested with BamHI and KpnI to excise the additional sgRNA cassette. The linearized backbone was blunt-ended by Klenow and subsequently ligated to re-circularize the plasmid. The pLenti-CRISPR-sgRNA(*Trp53*) plasmid was modified from pLenti-CRISPRv2 (Addgene #52961).[67] First, the puromycin resistance gene sequence was disrupted via insertion of stop codon oligos downstream of Cas9 using BamHI and BstEII digestion. The oligos used for cloning here were: BamHI-Stop-T: 5’-gatccTAATAGTTATAGg-3’; BstEII-Stop-B: 5’-gtgacCTATTACTATTAg-3’. Subsequently, the same *Trp53* sgRNA oligos mentioned above were cloned into this plasmid via Esp3I as originally designed.[67] The pBabe-KrasG12D plasmid was reconstructed from pBabe-Kras^G12D^-puro (Addgene #58902) by removing the puromycin resistance gene through digestion with HindIII-HF and BspDI, blunt-ended by Klenow and re-ligation to restore plasmid circularity. All target sequences and sgRNA oligonucleotides used are indicated in **STable 2**.

### Lentivirus production

The Lenti-CRISPR-sgRNA(*Trp53*), pBabe-*Kras^G12D^*, or Lenti-luciferase-P2A-Neo plasmid (Addgene #105621) was co-transfected with packaging plasmid psPAX2 and pMD2.G (both from PlasmidFactory GmbH & Co. KG, Bielefeld, Germany) into HEK293T cells by TransIT LT1 transfection reagent according to the manufacturer’s instructions (Mirus Bio, Madison, WI, USA). The supernatant containing lentivirus was harvested 2 days after transfection, filtered, and stored at-80°C.

### Genetic engineering of murine cholangiocyte organoids

Before transfection, cholangiocyte organoids were dissociated by incubating with 500 µL/well of Cell Recovery Solution (Corning, #354253) on ice for 20 min, followed by centrifugation (500×g, 5 min, 4°C). Pellets were treated with 0.25% trypsin at 37°C for 5-8 min with intermittent shaking, then mechanically dissociated by syringe. After quenching with cold DMEM and centrifugation, cells were resuspended in chol GF+ media with 10µM of Y-27632 (TargetMol, #TMO-T1725-50mg) and plated at 450 µL/well in a 48-well plate. For transfection, Lipofectamine 2000 (Invitrogen, # 11668019) and plasmid DNA were each diluted in OPTI-MEM, combined (1.5 µg DNA/reaction), and incubated at room temperature for 15 min. The DNA-Lipofectamine complexes were added dropwise to the cells. Plates were centrifuged at 600×g for 60 min at 32°C, then incubated at 37°C for 4 hours. Cells were collected, centrifuged (845×g, 5 min, 4°C), resuspended in 100–150 µL BME, and seeded into 2–3 wells of a 24-well plate. After 30 min, 500 µL of chol GF+ media was added.

To generate cholangiocyte organoids carrying *Trp53* deletion (chol-P), chol-WT organoids were transfected with pX458-sgRNA(*Trp53*) plasmid and treated with 7.5 μM Nutlin-3 (Biotrend #A4228-S) starting 2 days post-transfection for functional selection.

To generate cholangiocyte organoids carrying an additional *Kras^G12D^*mutation (chol-PK), cells were co-transfected with pX459-(BB) and sgRNA(*Kras*)-*Kras^G12D^*_HDR plasmids. The day after transfection, 1 µM of Alt-R™ HDR Enhancer V2 (IDT, #10007921) was added for 24 hours to promote homology-directed repair. On day 3, cells were treated with 1 μg/mL puromycin in chol GF+ media for 2 days to select for transfected cells. Beginning 3 days post-transfection, cells were switched to chol GF-media (lacking mEGF and hHGF) supplemented with 1 μM gefitinib for functional selection.

To generate KO lines, chol-PK were transfected with pX459 containing the respective sgRNA guides targeting *Plau*, *Lamc2*, *Prss22*, respectively. After 2-3 days of transient transfection, organoids were selected with 1-2 ug/µl of puromycin for 2-7 days, depending on selection efficiency. For all lines, chol GF-media were changed twice per week, and clones were picked approximately 14-21 days after transfection.

Primers used for genotyping the clones via amplification of genomic regions around the sgRNA target sites are indicated in **STable 2**. The PCR products were verified by Sanger sequencing (Microsynth AG, Balgach, Switzerland). Indel mutation frequencies were analysed using the Inference of CRISPR Edits (ICE) tool (https://ice.synthego.com/).[68] For the *Kras^G12D^* mutation in chol-PK, PCR amplicons were further confirmed by TA-cloning (PCR Cloning, NEB, #E1203S) and Sanger sequenced. Chol-PK were then transduced with the lentivirus carrying luciferase-P2A-Neo, and selected by 500 µg/ml G418 (ThermoFisher Scientific, #10131027) for at least 2 weeks.

### Genetic engineering of murine hepatocyte organoids

Before infection, hepatocyte organoids were dissociated using Accutase solution (PromoCell, # C-41310). Cells were plated at 100 μL/well in hep GF+ media supplemented with 10 μM Y-27632 and 8 μg/mL polybrene in a 48-well plate. For lentiviral infection, 100 μL of virus was added per well, and subsequent steps followed the same procedure used for cholangiocyte organoid transfection.

To generate *Trp53* deletion carrying hepatocyte organoids (hep-P), hep-WT organoids were transduced with Lenti-CRISPR-sgRNA(*Trp53*) virus and treated with 7.5 μM Nutlin-3 (Biotrend Cat# A4228-S) enriched hep GF+ media starting 2 days post-transfection for functional selection. The culture media was changed twice per week, and clones were picked between 14-20 days after infection.

To generate hepatocyte organoids carrying *Trp53* deletion and *Kras^G12D^* overexpression (hep-PK), hep-P organoids were transduced with pBabe-*Kras^G12D^* virus. The day after infection, cells were switched to hep GF-media supplemented with 1 μM gefitinib for functional selection. Media was changed twice per week, and initial clones were dissociated every 2 weeks. Pooled clones were maintained in the selection media for a total of 33 days post-infection. Hep-PK organoids were further transduced with the lentivirus carrying luciferase-P2A-Neo and selected by 500 µg/ml G418 (ThermoFisher Scientific, #10131027) for at least 2 weeks.

### Animal experiments

C57BL/6J mice were obtained from Janvier Labs and housed at the German Cancer Research Center (DKFZ) under specific pathogen-free (SPF) conditions. Mice were maintained in individually ventilated cages on a 12-hour light/dark cycle at 21–23 °C, with ad libitum access to food and water. All experiments were conducted in accordance with German law and approved by the Regierungspräsidium Karlsruhe (approval no. G34/22). For orthotopic tumorigenicity assays, adult mice of both sexes older than 8 weeks were used. Mice were anesthetized with isoflurane (0.5-1% with oxygen), a small midline laparotomy and intrahepatic implantation via 28-gauge syringe into the left liver lobe was performed. The injection site was then compressed with a cotton-top to avoid leakage or bleeding, and peritoneum and skin closed with absorbable suture (Ethicon, #VCP303H) and 9 mm clips (Fine Science Tools, #12022-09), respectively. Proper peri-and postoperative analgesia was applied with metamizole and lidocaine. For implantation, chol-PK organoids and their KO derivatives at passages 7-15 (1.5-2 × 10⁵ cells/implantation) and Hep-PK organoids at passages 8-13 (2-5 × 10⁵ cells/implantation) in 25 µl DMEM:BME mixture (1:1) were used.

The *in vivo* imaging System (IVIS) was used for tumor growth surveillance. To this end, the abdominal area of the mice was shaved and 150 mg/kg D-Luciferin (Promega, #E1605) injected intraperitoneally, and imaging performed under anesthesia with isoflurane (1–2 vol.% in oxygen).

### Histopathology and immunohistochemistry

For fixation of organoids, upon aspiration of culture medium, BME was removed by incubation with cell recovery solution for 15 min on ice (Corning, #354253). Then, organoids were washed with PBS, fixed using 4% paraformaldehyde, and then embedded into HistoGel (Thermo Scientific. #HG-4000-012) as per manufacturer’s instructions. Histogel blocks were fixed overnight in 10% neutral buffered formalin solution, washed with PBS and 70% ethanol, and embedded in paraffin. Serial sections of 5 µm thickness were generated and hematoxylin and eosin (H&E) staining was performed following routine histological protocols.

For immunohistochemical (IHC) staining, sections were deparaffinized, rehydrated, and subjected to heat-induced antigen retrieval with either citrate buffer at pH 6.0 (Vector, H-3300) for CK19 and HNF4α, or Tris-based buffer at pH 9.0 (Vector, H-3301) for CD44. After washing, endogenous peroxidase activity was quenched with 0.3% H₂O₂, and nonspecific binding was blocked using blocking solution (5% donkey serum (Biozol, #LIN-END9010) in 1% BSA/PBS containing 0.1% Tween-20) for 1 hour at room temperature. For mouse-derived primary antibodies, additional blocking was performed with a M.O.M. reagent (Vector, #MKB-2213-1) according to the manufacturer’s instructions. Sections were then incubated overnight at 4 °C with primary antibodies, followed by appropriate biotinylated secondary antibodies. Detailed information for primary and secondary antibodies are listed in **STable 3**. Detection was performed using the Vectastain Elite ABC-HRP Kit (Vector, #PK-7100) with diaminobenzidine (Vector, #SK4100) as chromogen. Nuclei were counterstained with Hematoxylin solution (Carl Roth, #T864.1), and slides were dehydrated and mounted with Vectamount mounting medium (Vector, #H-5700-60).

### Tissue mounting and immunostaining for multiplexed ion beam imaging (MIBI)

Formalin-fixed and paraffin-embedded tissue blocks were sectioned at 5 µm and mounted onto gold-coated conductive slides (Ionpath, #567001). To minimize batch effects, all samples were processed together in parallel and stained with common antibody master mixes, following established protocols.[69] Briefly, sections were deparaffinized and rehydrated as for IHC preparation, then subjected to antigen retrieval. Non-specific binding was blocked for 1 h at room temperature, followed by M.O.M. reagent (Vector, #MKB-2213-1) to prevent background from endogenous mouse IgG. Slides were then incubated overnight at 4 °C with a filtered primary antibody master mix for indirect staining. The following day, sections were washed and incubated with a filtered secondary antibody master mix for 1 h at room temperature. After washing, slides were incubated overnight at 4 °C with a filtered direct-staining antibody master mix. All antibodies used for MIBI staining are listed in **STable 4**. Following a final wash, sections were fixed in 2% (v/v) glutaraldehyde for 5 min at room temperature and dehydrated through a graded ethanol series. Prior to acquisition, stained slides were stored in a vacuum cabinet to prevent metal oxidation and preserve tissue integrity.

### Multiplexed imaging and data prepressing

H&E images of serial sections corresponding to the MIBI slides were used to identify tumor boundary regions for imaging. For each sample, one region was manually selected in DUEfiner, a customized interactive image viewer implemented in Python, and subdivided into 400 × 400 µm fields of view (FOVs). Each FOV was acquired at a dwell time of 0.5 ms using the “Coarse” preset, yielding a pixel size of 390 nm.

Images were extracted from mass spectrometry data using Toffy (https://github.com/angelolab/toffy). For each pixel, the intensity of a given mass 𝑚 was obtained by integrating ion counts over the range [𝑚 − 0.3, 𝑚]. To correct for systematic intensity gradients across FOVs — analogous to illumination correction in light and fluorescence microscopy —, all images were examined for such patterns. A gradient was defined as a consistent intensity variation present across channels and multiple FOVs. For each channel, the gradient was estimated by summing Gaussian-blurred versions of its FOVs, thereby reducing contributions from biological signals. Gradient magnitude scaled with channel mass, and a per-channel scaling factor was derived by linear regression between a proxy of gradient magnitude and channel mass. To further suppress biological signal, a two-dimensional correction map was generated by summing the scaled gradients across all channels. Correction was applied by dividing each channel by this map, adjusted for the channel-specific gradient magnitude across all FOVs.

Gradient-corrected images were subsequently processed with Rosetta compensation to reduce cross-channel contamination, as previously described.[70] The mass and relative abundance (coefficient) of interfering species in the source reagent are well characterized,[71] enabling estimation of their contribution by scaling the source channel intensity before subtracting it from the target channel. A coefficient matrix for all source–target channel pairs is available at https://doi.org/10.1101/2025.01.26.634557. To remove gold-signal-based background, bare slide regions were identified and excluded based on elevated gold (Au) ion signal from the conductive slide coating. Pixel intensities in the Au channel were normalized to the 99.9th percentile and arcsinh-transformed (cofactor = 100). The transformed intensity distribution was modeled with a double-Gaussian fit to distinguish gold-negative and gold-positive populations. The intersection of the two Gaussian components was defined as the threshold for gold-positive pixels. This threshold was then back-transformed into raw intensity space and applied to each FOV to generate binary masks of gold-positive pixels. These masks were subsequently used to exclude bare slide regions from downstream analysis.

### Image-based single cell data quantification and filtering

FOV tiles were stitched into whole-region images. Cells were segmented in the stitched images using the Cellpose TN model,[72] re-trained on unpublished MIBI datasets comprising 335 cells from human normal liver, normal lung, colorectal cancer, and lung cancer tissues. The model was applied with a cell diameter of 30 pixels, a flow threshold of 0.7, and a cell probability threshold of –5. Cell masks were filtered based on size, nuclear intensity, and gold-pixel content to retain only high-confidence single cells. Masks smaller than 100 pixels were removed as over-segmented. For retained cells, features including spatial location in the whole-region image, cell size, and cell-size–normalized (mean) channel intensities were calculated. Histone H3 (HH3) intensities were either summed or averaged per cell, percentile-scaled, arcsinh-transformed, and thresholded using the intersection of a double-Gaussian fit to identify nuclei-negative populations. Cells below this threshold were flagged as out-of-plane. Similarly, cells with >50% of their area overlapping bare-slide pixels were flagged as incomplete. All flagged cells were excluded from downstream analysis.

### Data annotation and tissue zone identification

To generate robust and accurate cell labels, we employed an iterative clustering strategy with human-in-the-loop curation across multiple levels, defined by the hierarchy of lineage markers (**Figure 3A**). At each level, FlowSOM [73] was used to cluster cells into self-organizing map (SOM) nodes based on lineage markers relevant for that level. Nodes were then merged to align with defined cell lineages according to marker positivity and visual inspection. Nodes with ambiguous marker profiles were re-clustered and annotated; cells that remained ambiguous were labeled as “other.” Importantly, at each level, cells belonging to different higher-level lineages were classified separately to maintain lineage-specific resolution.

At level 1, cells were broadly classified into hepatocytes, cancer, endothelial, fibroblasts, and immune cells. At level 2, fibroblasts were further classified according to Vimentin (Vim), αSMA, COL1a1, and PDGFRb positivity, while immune cells were subdivided into CD3⁺, CD20⁺, CD11c⁺, CD11b⁺, and CD11b⁺CD11c⁺ populations. At level 3, CD3⁺ cells were further divided into CD4⁺, CD8⁺, and CD4⁺FOXP3⁺ subsets, and CD11b⁺ cells into F4/80ᵐᵉᵈ, Ly6G⁺, and F4/80^hi^ subsets. At level 4, F4/80^hi^ macrophages were split into CD163⁻ and CD163⁺ populations. Immune cells with ambiguous finer lineage profiles were marked as “other immune.” All annotation steps were performed using our custom viewer, UELer (https://github.com/HartmannLab/UELer).

Tissues were divided into three zones: adjacent normal, tumor periphery, and tumor core. The tumor border was manually marked as the interface where hepatocytes confronted tumor cells, using tissue images with level 1 markers rendered in UELer. Relative to this border, the hepatocyte side was defined as adjacent normal, while the opposite side was defined as tumor. The tumor region was further subdivided into periphery and core based on distance from the border, using a cutoff of 150 µm.

Pixel-level annotations were performed using Ilastik [74] to identify SMA⁺ regions and stromal regions. The Ilastik workflow consisted of two phases: labeling and prediction. In the labeling phase, sparse labels corresponding to user-defined pixel classes were manually created. In the prediction phase, the Ilastik pixel classifier trained on these labels was applied to the entire dataset to assign class annotations per pixel.

To identify SMA⁺ regions, image stacks of αSMA, COL1a1, PDGFRb, PanCK, and CD44 were used as input. The features “Color/Intensity”, “Difference of Gaussians” and “Hessian of Gaussian Eigenvalues,” each Gaussian smoothed with σ = 3.5 and 10 pixels, were included for training the predictive model. Three annotation classes were defined: “SMA⁺“, “SMA⁻“, and “other“. SMA⁺ labels marked SMA⁺ pixels within tumor regions, SMA⁻ labels marked SMA⁻ pixels within tumor regions, and “other” labels marked pixels outside tumor regions (i.e. normal liver or luminal regions). The additional markers (COL1a1, PDGFRb, PanCK, CD44) were not used during labeling but served as supportive channels during prediction.

To identify stromal regions, image stacks of ATP5A, LDH, G6PD, HNF4a, and HH3 were used as input. The first three markers are cytoplasmic metabolic regulators and were used as indicators of cell bodies, while HH3 and HNF4a served as nuclear indicators. The features “Color/Intensity,” “Laplacian of Gaussian,” “Gaussian Gradient Magnitude,” “Difference of Gaussians,” “Structure Tensor Eigenvalues,” and “Hessian of Gaussian Eigenvalues,” each Gaussian smoothed with σ = 1 and 7 pixels, were included for training the predictive model. Four annotation classes were defined: “cell”, “non-cell”, “cyst & vessel”, and “outside tissue.” Pixels annotated as “cell” by the trained Ilastik model were defined as cell masks. To exclude vessels and cyst-like structures, tissue region masks encompassing cells and small extracellular gaps were generated by applying morphological opening and closing operations (kernel sizes 2 × 2 and 9 × 9, respectively) to fourfold downsampled cell masks. The remaining large voids were identified as vessels or cyst-like structures and excluded from downstream analysis. The extracellular space ratio within tumor tissue was calculated as the total area of remaining extracellular gaps divided by the total tumor tissue area for each sample.

### MIBI statistical analysis

To account for intra-tissue heterogeneity, most comparisons were performed at the tissue-patch level. Tissue patches were assigned according to the nearest tumor border segments. Depending on the hypothesis being tested, each patch was further subdivided into the three tissue zones (see Tissue zone identification) or into SMA proximity zones according to the SMA⁺ masks (see Pixel annotations).

Cell densities were calculated as the counts of a specified cell type divided by the total area of all cells within the specified region (e.g., tumor periphery per patch). Patch-based cell densities between groups were compared using two-sample t-tests. Densities between SMA proximity zones were compared using paired t-tests. Single-cell–level comparisons were performed using the Mann–Whitney U test. All significance values were determined from p-values corrected for multiple testing using the false discovery rate (FDR) method, applied across comparisons within the same figure panel.

### RNA sequencing and integrative data analysis

RNA was isolated using the RNeasy Plus Mini kit (Qiagen, #74134), and RNA integrity was analyzed by a 2100 Bioanalyzer using the RNA 600 Nano Kit (Agilent technologies) according to the manufacturer’s protocol. Sequencing libraries were prepared by the Genomics and Proteomics Core Facility (DKFZ, Heidelberg, Germany) from total RNA using the Illumina TruSeq Stranded Total RNA Library Prep Kit according to the manufacturer’s instructions. For sequencing, multiplexed libraries were sequenced in a paired-end setting (100 bp) on an Illumina NovaSeq 6000 sequencer.

Data was analyzed with the DKFZ Omics IT and Data Management Core Facility (ODCF) RNAseq workflow (version 4.0.0). Reads were aligned to the mouse genome mm10 using the STAR aligner version 2.5.3a. The raw counts were obtained using the featureCounts tool.[75] Differential expression was performed using DESeq2 version 1.44.0 in R 4.4.3.[76] We fit a two-way negative binomial generalized linear model (∼ lineage + strain + genotype + lineage:genotype) that adjusts for strain and includes a lineage×genotype interaction, allowing tests of lineage differences within each genotype, genotype differences within each lineage, and whether genotype effects differ by lineage (interaction). Genes with adjusted P values less than 0.05 and absolute log2 fold changes greater than 1 were identified as differentially expressed genes (DEGs) for each comparison.

Genes ranked by the DESeq2 Wald statistic were subjected to gene set enrichment analysis (GSEA) using the fgsea R package (v1.32.4).[77] Mouse Hallmark gene sets were retrieved using the msigdbr R package (v25.1.1) from the Molecular Signatures Database (MSigDB v2025.1, Mus musculus).[78,79] Enrichment calculations applied default fgsea settings with a gene-set size range of 3–500 genes. Pathways with FDR-adjusted p < 0.05 were regarded as significant, and normalized enrichment scores (NES) and adjusted p values were reported.

Per-sample lineage fidelity was estimated using gene set variation analysis (GSVA) implemented in the GSVA R package (v2.0.7).[80] Marker gene sets representing hepatocyte and cholangiocyte identities were derived from PanglaoDB cell-type annotations (accessed May 2025).[81] Lineage fidelity was defined as the difference between the GSVA scores for cholangiocyte and hepatocyte marker sets, with positive values indicating a cholangiocyte bias and negative values a hepatocyte bias. Hepatocyte markers (*Alb, Hnf4a, Asgr1, Afp, Foxa3, Apoa1, Apoa2, Ttr, Apom, C4b*, and *Serpina1*) and cholangiocyte markers (*Krt19, Mmp7, Krt7, Elf3, Sox9, Hnf1b, Aqp1*, and *Cftr*) were both obtained from PanglaoDB and used for GSVA.

The proportion of gene expression variance explained by experimental factors was estimated by fitting a linear mixed model using the variancePartition R package (v1.40.0).[82] The model included lineage, oncogenic effect, their interaction, and mouse strain as predictors (∼ lineage + genotype + lineage:genotype + strain), applied to variance-stabilized counts from DESeq2. The analysis was restricted to DEGs, defined as the union across the four contrasts (Chol-PK vs Chol-WT, Chol-PK vs Hep-PK, Chol-WT vs Hep-WT, and Hep-PK vs Hep-WT).

Secreted genes were obtained from the Human Protein Atlas database (https://www.proteinatlas.org/, accessed July 2024). Human-to-mouse gene orthologs were determined using the Mouse Genome Informatics (MGI) database (https://www.informatics.jax.org/, accessed in January 2025).

### Transcriptome comparison from public databases

Human gene expression data were obtained from The Cancer Genome Atlas (TCGA; https://www.cancer.gov/tcga), Cancer Cell Line Encyclopedia (CCLE), and GSE26566.[35,36] For the TCGA data, preprocessed RNA-seq data (STAR-aligned) for primary intrahepatic cholangiocarcinoma (TCGA-CHOL, n = 31) and primary hepatocellular carcinoma (TCGA-LIHC, n = 363) were used. To analyse CCLE data, DepMap 24Q4 and batch-corrected TPM values for CCA (n = 37) and HCC (n = 23) cell lines were used. GSE26566 consists of human CCA samples (n = 104) and adjacent non-tumor liver tissues (n = 59) with background-corrected and normalized microarray expression profiles.[36] Differential gene expression analyses were performed separately for each dataset using the limma package (v3.56.2).[83] Log₂-transformed values (pseudocount + 1) were used as input. Empirical Bayes moderation was applied, and p-values were adjusted for multiple testing using the Benjamini–Hochberg procedure.[84]

### Concentrated conditioned media (CCM) collection

Organoids were seeded in 12-well plates, with three 50 μL drops of BME per well. Once the organoids reached a high density, they were washed twice with prewarmed PBS, followed by incubation in 1 mL of DMEM per well for 16-24 hours to induce starvation. After starvation, conditioned media was collected, centrifuged at 1,000 × g for 5 minutes, and filtered through a 45 μm syringe filter to remove debris. The resulting supernatant was then concentrated ∼10-fold using a 10 kDa molecular weight cut-off filter (Vivaspin 6, Sartorius Lab, #VS0601) and centrifuged at 3,000 × g for 30 minutes to obtain CCM.

### LC-MS/MS analysis using Orbitrap Exploris 480 mass spectrometer

For the secretome analysis, CCM was collected and supplemented with a protease inhibitor cocktail (MedChemExpress, #HY-K0011) and stored at –80 °C until further processing. For the proteome analysis, organoids were collected after BME dissociation with cell recovery solution, mechanically disrupted through a 25G needle, and subsequently lysed in RIPA buffer (Thermo Fisher Scientific, #89900) supplemented with a protease inhibitor cocktail. Protein concentrations were determined using the BCA Protein Assay Kit (ThermoFisher Scientific, #23225) according to the manufacturer’s instructions. A total protein amount of 10 µg from either CCM or organoid lysates was digested with trypsin using an AssayMAP Bravo liquid handling system (Agilent technologies) running the autoSP3 protocol according to Müller et al.[85]

LC-MS/MS analyses were performed using either an Ultimate 3000 UPLC system (murine samples) or a Vanquish Neo UPLC system (human samples) (both Thermo Fisher Scientific), each directly coupled to an Orbitrap Exploris 480 mass spectrometer. The total run time was 120 min for murine samples (CCMs and organoid lysates) and human organoid lysates, or 60 mins for human CCMs. Peptides were online desalted using a trapping cartridge (Acclaim PepMap300 C18, 5 µm, 300 Å wide pore; Thermo Fisher Scientific): 30 µl/min flow of 0.05% TFA in water was used for 3 min (murine samples) or with a loading volume of 60 µl (human samples). Separation was carried out on a nanoEase MZ Peptide analytical column (300 Å, 1.7 µm, 75 µm × 200 mm, Waters) using a multistep gradient at 300 nl/min with solvent A (0.1% formic acid in water) and solvent B (0.1% formic acid in acetonitrile). For murine samples, the concentration of B was linearly ramped from 4% to 30% for 102 min, followed by a quick ramp to 76%, after two minutes the concentration of B was lowered to 2% and a 10 min equilibration step appended. For human samples, the concentration of B was linearly ramped from 5% to 30% for either 105 min (120 min method) or 46 min (60 min method), followed by a quick ramp to 80%, after four minutes the concentration of B was lowered to 2% and a three-column volumes equilibration appended. Eluting peptides were analyzed using data-independent acquisition (DIA) mode.

For murine samples, a MS1 scan at 120k resolution (380-1400 m/z, 300% AGC target, 45 ms maxIT) was followed by 47 overlapping (1 Da) DIA windows of variable width covering m/z-range of 400-1000. For human samples, the MS1 scan was recorded at 120k resolution (380-1400 m/z, 300% AGC target, 45 ms maxIT). The following DIA windows (20 for 60 min method and 40 for 120 min method) were of variable width ensuring a 1 Da overlap covering the m/z-range of 400-1000 m/z.

DIA windows were acquired using a HCD collision energy of 28% and were recorded at a resolution of 30k with an AGC target of 1000% and a maxIT of 54 ms. Each sample was followed by a wash run (40 min) to minimize carry-over between samples. Instrument performance throughout the course of the measurement was monitored by regular (approx. one per 48 hours) injections of a standard sample and an in-house shiny application.

### LC-MS/MS data analysis

All sample handling (sample preparation and LC-MS/MS analysis) was conducted in a block randomization design to minimize batch effects.[86]

For murine samples, analysis of DIA RAW files was performed with Spectronaut (Biognosys, version 17.1.221229.55965) in directDIA+ (deep) library-free mode.[87] Default settings were applied with the following adaptions. Within the Pulsar Search in Result Filters the m/z Max was set to 1800 and the m/z Min to 300. Separate LFQ normalization was performed via different fractions for conditioned media and cell lysates. The analysis was performed once with and once without normalization. The data was searched against the mouse proteome from Uniprot (mouse reference database with one protein sequence per gene, containing 21,957 unique entries from 3rd of Mai 2023) and the contaminants FASTA from MaxQuant (246 unique entries from 22nd of December 2022).

For human samples, analysis of DIA RAW files was performed with Spectronaut (Biognosys, version 19.1.240724.62635) in directDIA+ (deep) library-free mode.[87] Default settings were applied with the following adaptions. Within DIA Analysis under Identification the Precursor PEP Cutoff was set to 0.01, the Protein Qvalue Cutoff (Run) set to 0.01 and the Protein PEP Cutoff set to 0.01. In

Quantification the Proteotypicity Filter was set to Only Protein Group Specific, the Protein LFQ Method was set to MaxLFQ and the Quantification window was set to Not Synchronized (SN 17). Separate LFQ normalization was performed via different fractions for CCMs and oorganoid lysates. The analysis was performed once with and once without normalization. The data was searched against the human proteome from Uniprot (human reference database with one protein sequence per gene, containing 20,597 unique entries from 9th of February 2024) and the contaminants FASTA from MaxQuant (246 unique entries from twenty-second of December 2022). Conditions were included in the setup.

### LC-MS/MS statistical analysis

Statistical analyses were performed separately for murine and human datasets using log₂-transformed protein quantity (LFQ)values. For murine samples and human organoid lysates, the statistical analysis was performed with the fraction-specific LFQ normalized quantity values. For human CCMs, the statistical analysis was performed with the non-normalized quantity values, as the assumption that most protein abundances remain constant between conditions could not be upheld. This approach ensured consistent protein grouping across sample preparations. The following steps were performed per statistical contrast and not across the whole data matrix. Protein groups with valid values in 70% of the samples of at least one condition were used for statistics. For human CCM, the statistical analysis was performed without additional normalization. For human organoid lysates quantile normalization was applied in addition to LFQ normalization to improve comparability across samples.[88] Missing value imputation followed a two-step strategy adapted from Perseus.[89] When values were completely absent in one condition, they were imputed with random values drawn from an intensity distribution that was downshifted by 2.2 standard deviations and narrowed to 0.3 standard deviations. In cases where missing values were not fully absent in one condition, imputation was performed using the R package missForest, a non-parametric method suitable for mixed-type data.[90]

Differential expression analysis was conducted using the R package limma,[83] with model parameters based on the “Two Group” sections of the user guide. The empirical Bayes moderation was adapted for proteomics data by setting the trend and robust options to TRUE, as recommended by the authors. P-values were adjusted for multiple testing using the Benjamini–Hochberg method.[84]

#### HSC treatment experiments

Immortalized murine hepatic stellate cells SV40 (mHSCs, Creative Bioarray, CSC-I9351L) were maintained in DMEM containing 10% FBS and 1% PenStrep. Before stimulation, cells were seeded in a 24-well plate with 50,000 cells per well or in a 96-well plate with 30,000 cells per well, and cultured overnight in DMEM containing 2% FBS and 1% PenStrep. For stimulation, the old media was removed, and the cells were subsequently treated with DMEM containing 5% of the indicated CCMs and a final concentration of 2% FBS. Following a 24-hour stimulation period, cells were harvested.

For antibody blocking, 30μg/ml of indicated antibodies were preincubated with 5% chol-PK CCM in DMEM (supplemented to a final concentration of 2% FBS) for 2 hours at RT. The antibodies used in this study were: Lamc2 (Proteintech, #19698-1-AP), uPA (Proteintech, #17968-1-AP), BSSP4 (ThermoFisher Scientific, #PA5-47245), and rabbit-IgG (Millipore, #pp64).

### RNA extraction and quantification by real-time PCR

RNA from mHSCs and liver tumors was isolated using the Direct-zol RNA microprep kit (Zymo Research, #R2062). Complementary DNA was prepared using Iscript cDNA Synthesis Kit (Bio-Rad, #1708891) and selected genes quantified by real-time PCR using Luna Universal qPCR Master Mix (NEB, #M3003E) run on a CFX96 real-time PCR machine (Bio-Rad). Relative gene expression was calculated by the 2^-ΔCt^ method. Target gene expression was normalized to the indicated housekeeping genes. Primers used in this study are listed in **STable 5**.

### Cell scratch assay

A total of 30,000 mHSCs in 100 μL medium were seeded per well in 96-well plates and incubated overnight to allow monolayer formation. The cell monolayer was scratched using the IncuCyte WoundMaker (Sartorius, Göttingen, Germany) to generate standardized wounds of 700–800 μm width across all wells, following the manufacturer’s instructions. Detached cells by scratching were removed by washing twice with 100 μL DMEM per well. The remaining adherent cells were treated with 5% of chol-PK CCM and recombinant proteins with the indicated concentration in 100 μL of 2% FBS-containing medium. The recombinant protein for mouse PLAU/uPA (MedChemExpress, HY-P78017) and mouse PRSS22/BSSP4 (Boster Bio, PROTQ9ER10) were used. Plates were placed in the IncuCyte S3 Live-Cell Analysis System (Sartorius), and images were captured every hour for up to 26 hours. Relative wound density was quantified using the built-in IncuCyte 2021C software. All conditions were evaluated in 2 – 3 independent experiments with more than 4 replicates each.

### Software

Data, unless indicated otherwise, were analysed and plotted using QuPath, Living Image (IVIS) and GraphPad Prism 10. Figure panels were generated by Affinity Designer 2. For the creation of some illustration panels, icons from BioRender.com were used.

## Extended Data

**Supplementary Figures**

**SFigure 1.**
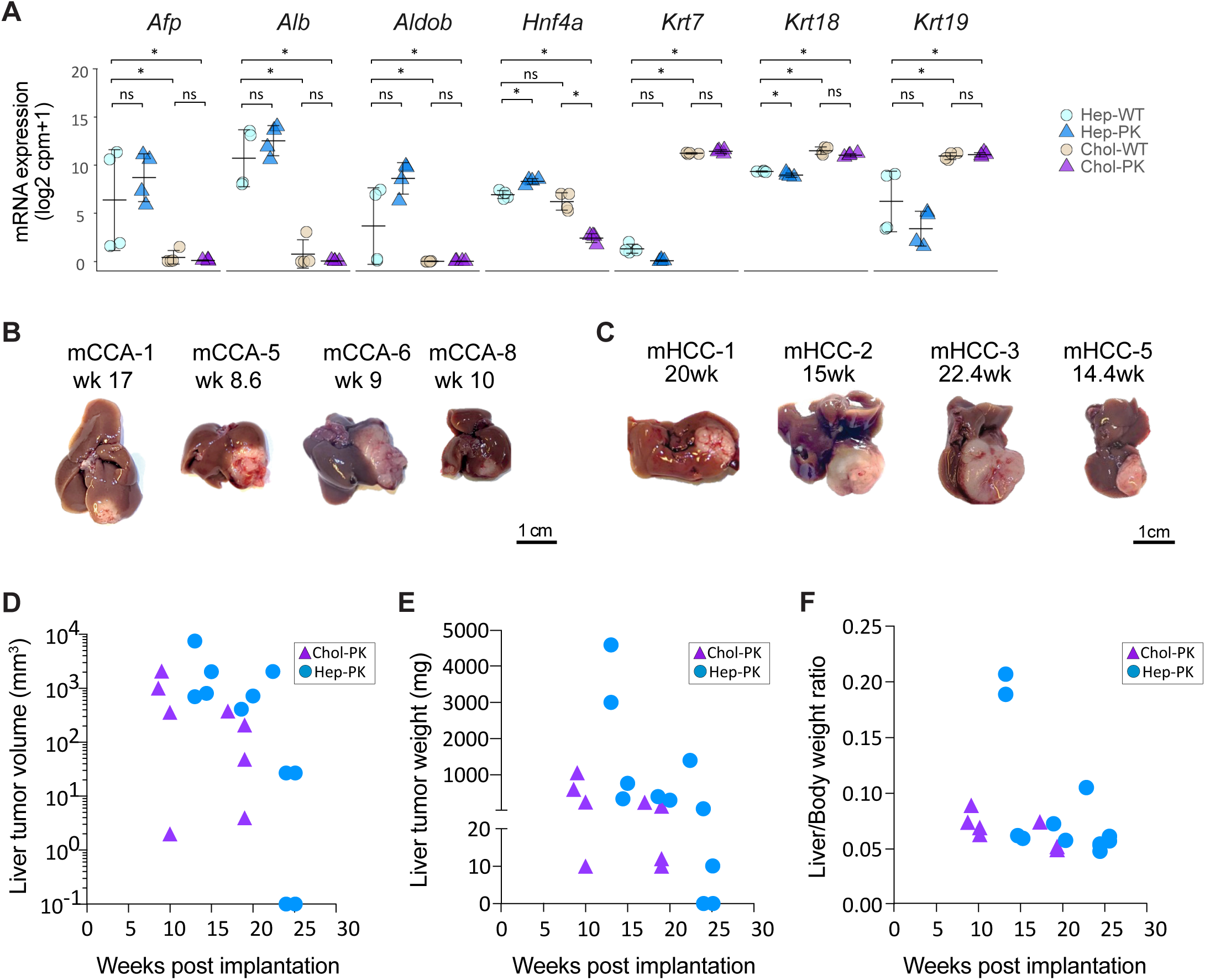
Cholangiocyte-and hepatocyte-organoid-derived liver tumors. (A) Scatter plot of mRNA expression of hepatocyte markers (*Afp, Alb, Adob*, and *Hnf4a*) and cholangiocyte markers (*Krt7, Krt18*, and *Krt19*) in wildtype (WT) and Trp53^del^ and Kras^G12D^-mutated (PK) hep-and chol-orgs. N=4 per group; * indicates *p* ≤ 0.05. (B-C) Macroscopic images of liver tumors obtained upon implantation of chol-PK organoids (B, as mCCAs) and hep-PK organoids (C, as mHCCs) and indicating harvest time point in weeks (wk) after implantation. (D-F) Liver tumor volume (D), liver tumor weight (E), and liver/body weight ratio (F) for both chol-PK and hep-PK derived liver tumors plotted against harvest time point as weeks post implantation. Data points represent individual mice.

**SFigure 2.**
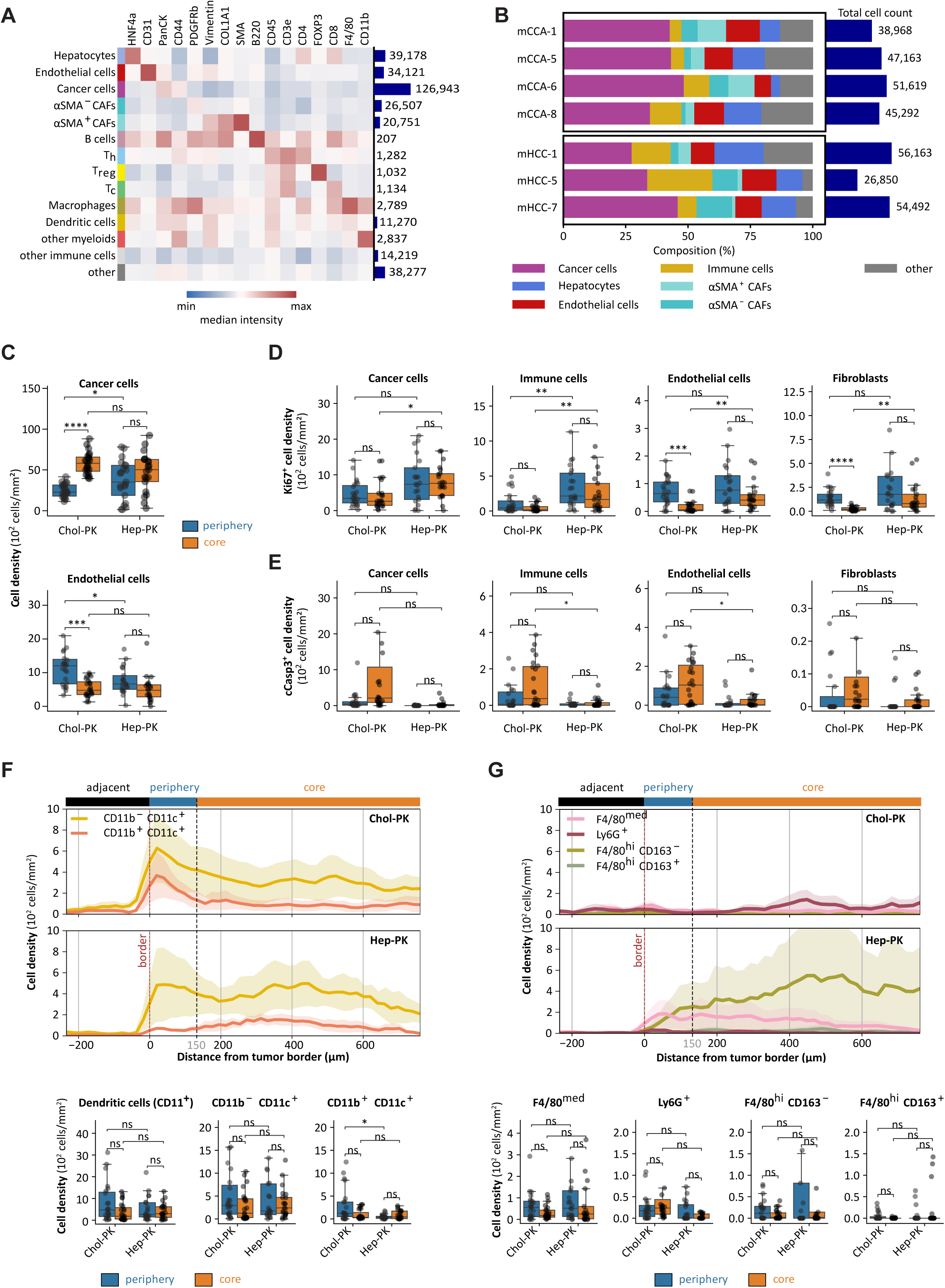
Spatial and phenotypic TME profiling in iCCA and HCC. (A) Heatmap showing z-scored median expression intensities of markers across major annotated cell types from 4 chol-PK CCAs and 3 hep-PK HCCs. Total cell count per cell type is indicated to the right of the heatmap. (B) Cellular composition of each tumor sample, highlighting proportions of immune cells, cancer cells, hepatocytes, endothelial cells, and αSMA⁺ and αSMA⁻ CAFs. Total cell count per sample is indicated to the right. (C) Box plot depicting cancer cell and endothelial cell densities separated by tumor periphery and core regions. (D) Box plot depicting proliferating (Ki-67⁺) cells across tumor compartments (periphery versus core) for cancer cells, immune cells, endothelial cells, and fibroblasts. (E) Box plot depicting apoptotic (cCasp3⁺) cells across tumor compartments (periphery versus core) for cancer cells, immune cells, endothelial cells, and fibroblasts. (F) Spatial distribution measured as cell density along an orthogonal axis of the tumor (top) and box plots (below) depicting CD11c⁺ dendritic cells, including CD11b⁺CD11c⁻ and CD11b⁻CD11c⁺ densities across tumor periphery and core regions. (G) Spatial distribution measured as cell density along an orthogonal axis of the tumor (top) and box plots (below) depicting other myeloid cell subtypes, including macrophage subsets (F4/80^hi^CD163⁺ and F4/80^hi^CD163⁻), F4/80^med^ and Ly6G⁺ cells densities across tumor periphery and core regions. In (C-G), each data point represents one tissue patch. Statistical significance was defined as: *p* < 0.05 (*), *p* < 0.01 (**), *p* < 0.001 (***), and *p* < 0.0001 (****).

**SFigure 3.**
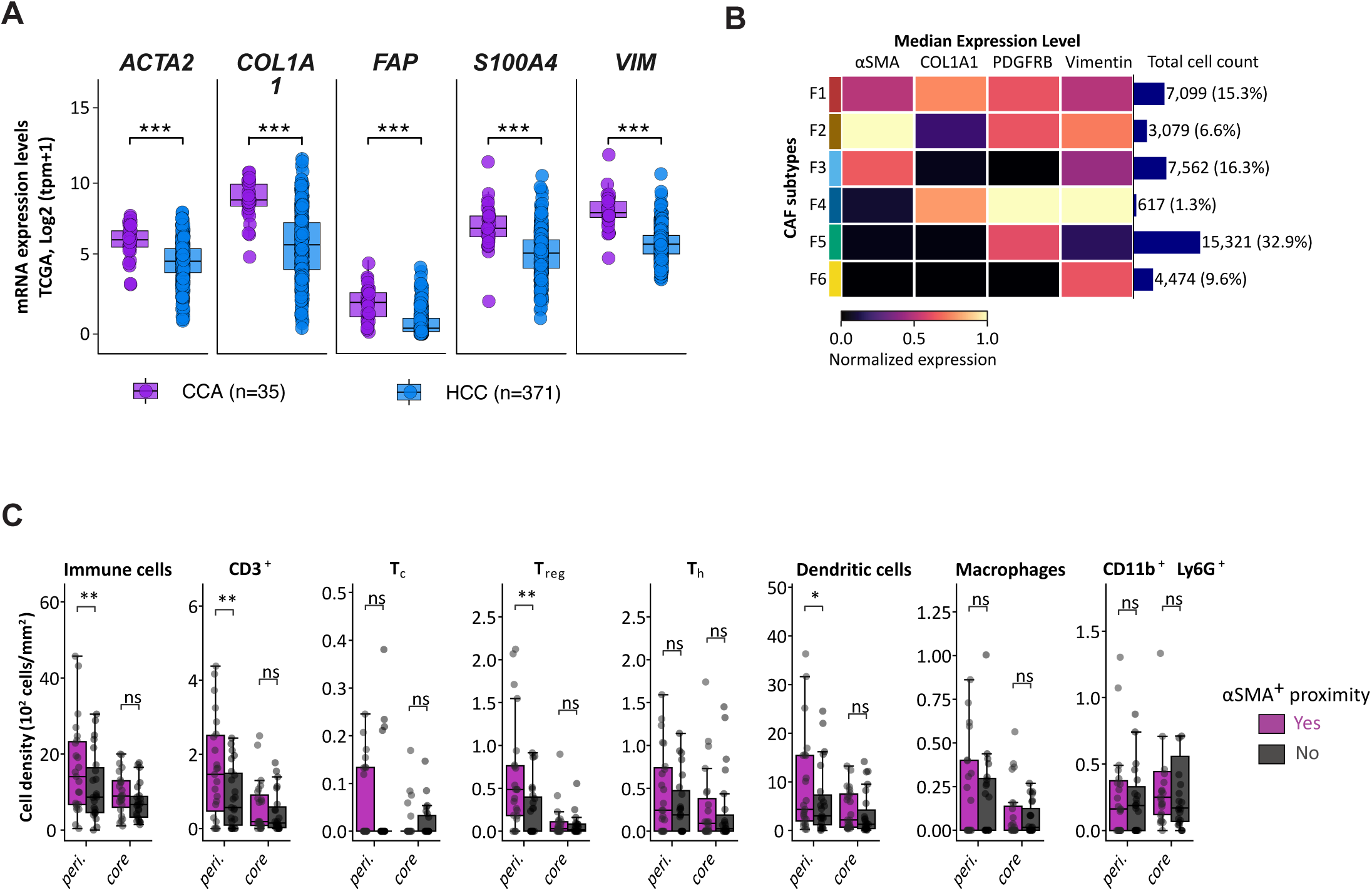
Characterization of CAF Distribution and Co-Localization with Immune Cells in iCCA. (A) Messenger RNA expression of 5 cancer-associated fibroblast (CAF)–related genes (*ACTA2, COL1A1, FAP, S100A4*, and *VIM*) in CCA and HCC from TCGA data set. Each data point represents one tumor sample. (B) Heatmap of normalized median expression levels of CAF markers (SMA, COL1A1, PDGFRβ, and Vimentin) across six fibroblast clusters (F1–F6). Total cell count and proportion of each subtype are listed on the right. (C) Density of immune cells in proximity (<20 μm) to αSMA⁺ CAFs within the periphery and core of chol-PK tumors. Each data point represents one tissue patch. Statistical significance was defined as: *p* < 0.05 (*), *p* < 0.01 (**), *p* < 0.001 (***), and *p* < 0.0001 (****).

**SFigure 4.**
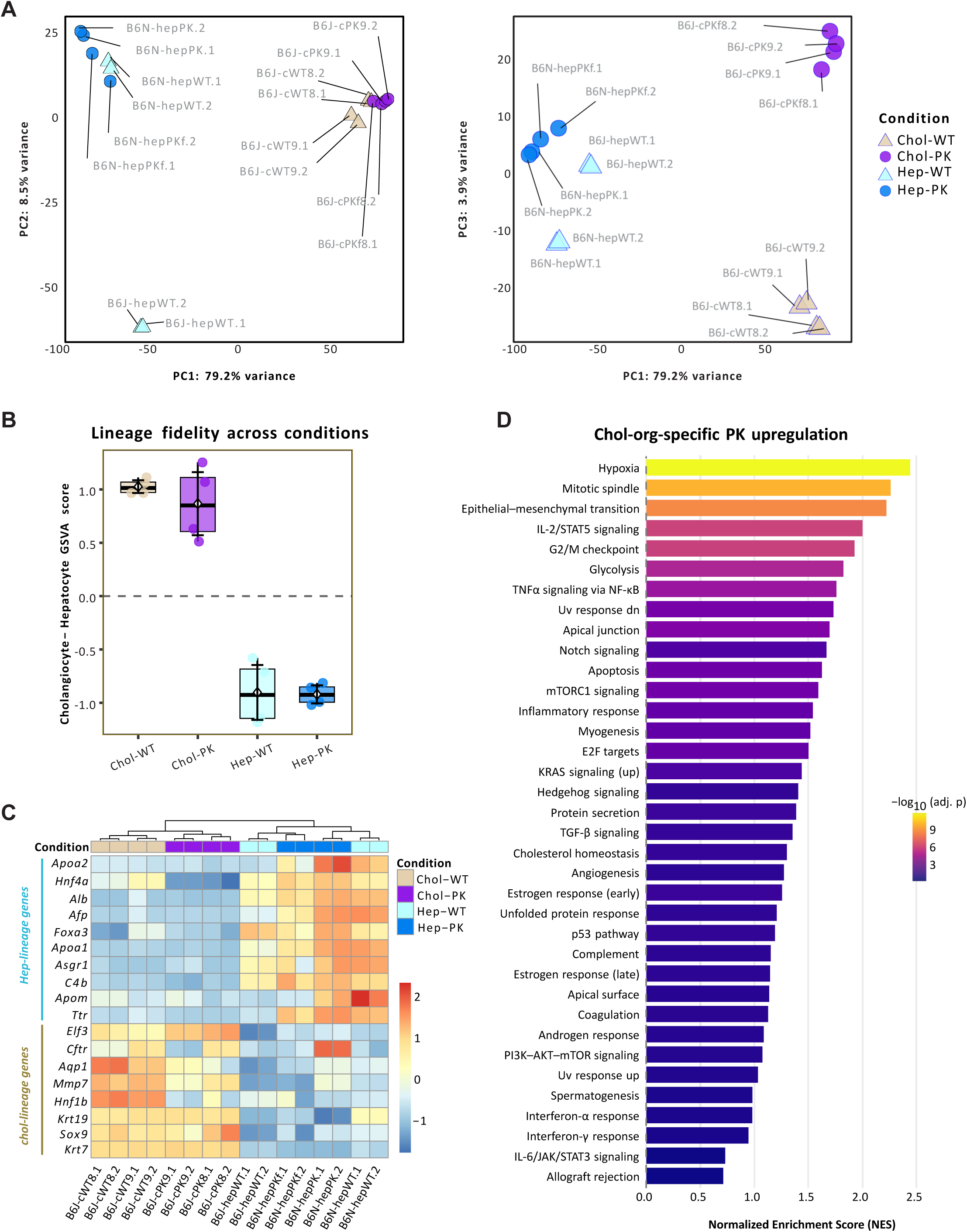
Extended Organoid Transcriptome Analysis. (A) Principal component analysis (PCA) of transcriptomes from wildtype and transformed murine hep-and chol-orgs with each individual sample labelled. Shown are PC1 (79.2% variance) versus PC2 (8.5%), which separate samples primarily by cell type and mouse-strain differences (C57BL/6J vs C57BL/6N), and PC1 versus PC3 as in Fig. 5B. (B) Box plot depicting lineage fidelity per sample computed as the GSVA score difference (cholangiocyte − hepatocyte), grouped by condition (chol-WT, chol-PK, hep-WT, hep-PK). Positive values indicate cholangiocyte bias, negative values hepatocyte bias, 0 marks absence of bias. Marker gene sets were taken from PanglaoDB (see Methods for details). (C) Heatmap of normalized expression for PanglaoDB-curated hepatocyte and cholangiocyte marker genes across organoid samples. Samples are hierarchically clustered and annotated by condition. The expression patterns validate the expected lineage assignment. (D) Gene set enrichment analysis (GSEA) of hallmark pathways specifically enriched in the lineage-dependent PK effect. We modelled gene expression with a linear model as a function of cell type, oncogenic effect (PK vs WT), and their interaction. Genes were ranked by the interaction test statistic, which quantifies the strength of the PK effect difference between lineages. All the significant pathways are shown.

**SFigure 5.**
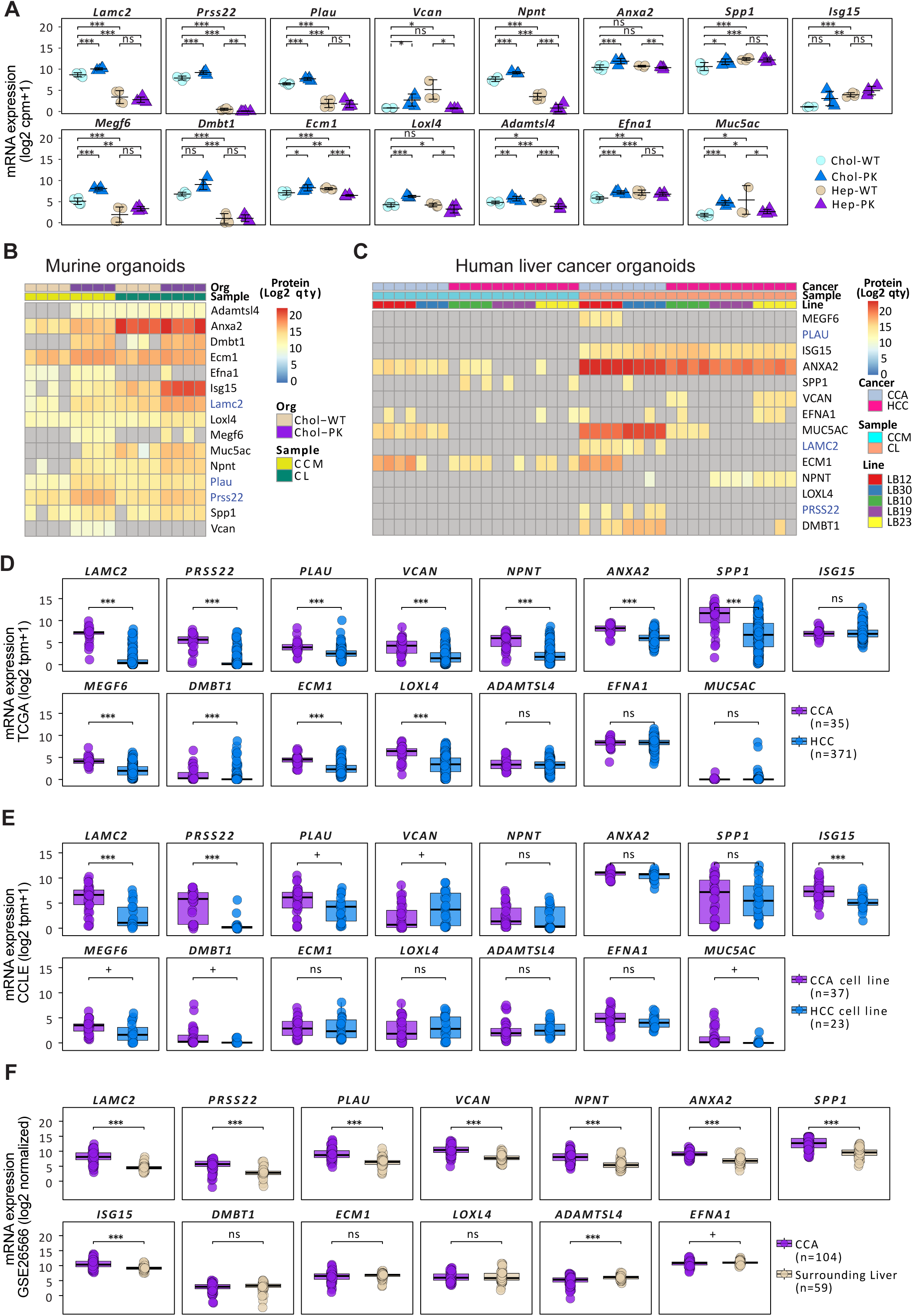
Multi-omics analysis from murine and human liver cancer organoids and public liver cancer databases. (A) Scatter plot of mRNA expression of 15 selected genes identified by the multi-omics analysis as depicted in Figure 6B. Organoid groups as indicated. (B-C) Heatmap of protein abundance (log₂ quantity) for the 15 selected proteins in concentrated conditioned medium (CCM) and cell lysate (CL) from murine chol-orgs (B) and patient-derived CCA and HCC organoids (C). (D-F) Box plots depicting mRNA expression of 15 selected genes in CCA and HCC from TCGA (D) and CCLE (E) data sets, and 13 available genes in CCA and non-tumorous liver from GSE26566 (F) data set. Each data point represents one sample. Statistical significance was defined as: *p* < 0.05 (*), *p* < 0.01 (**), *p* < 0.001 (***), and *p* < 0.0001 (****).

**SFigure 6.**
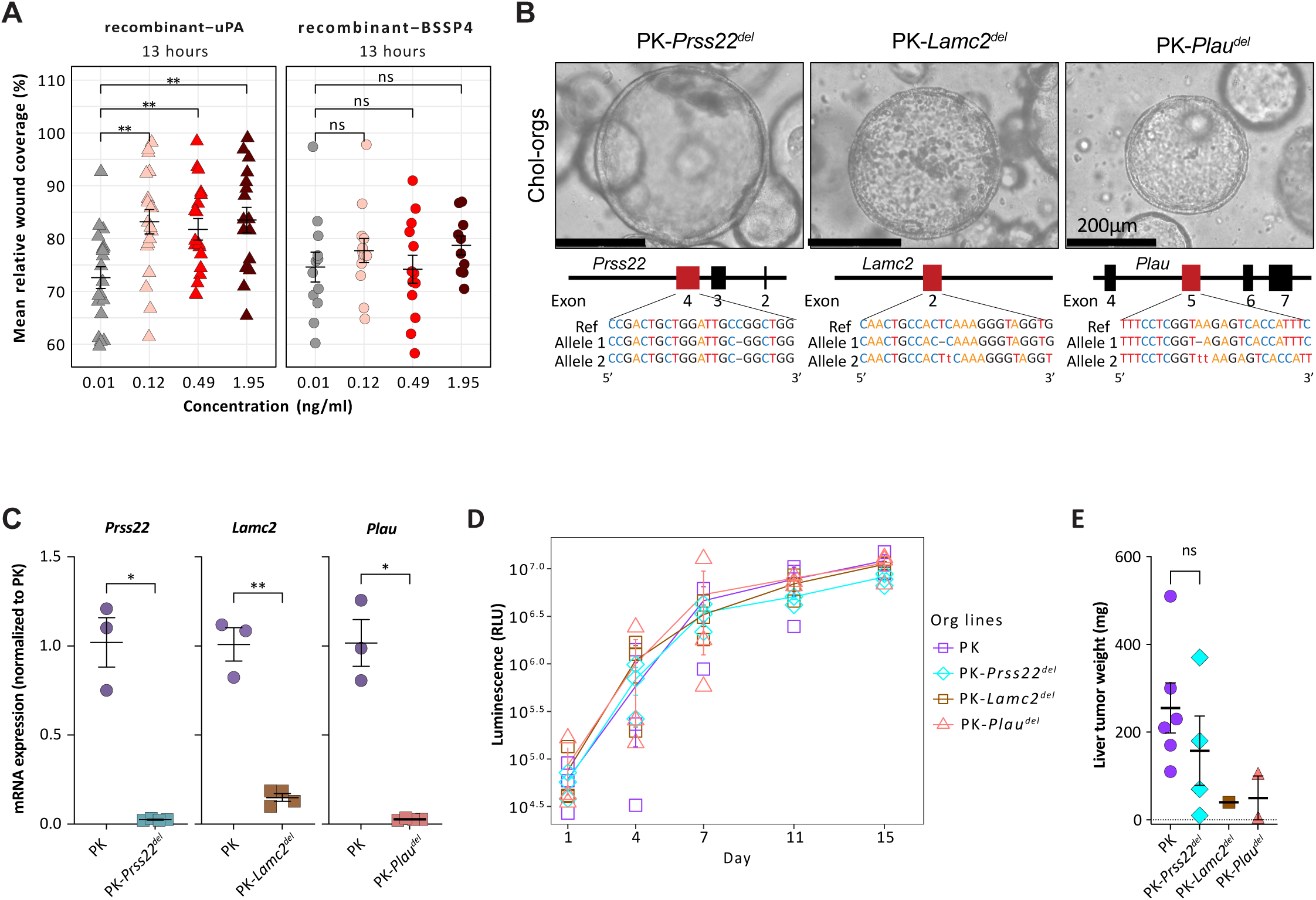
Characterization of Prss22^del^, Plau^del^ and Lamc2^del^ chol-PK organoid lines. (A) Scatter plot of scratch assay with mHSC depicting percentage of mean relative wound coverage at time point 13 hours with treatment of recombinant uPA or BSSP4 at the indicated concentrations (n≥10 from 3 experiments). (B) Bright-field microscopic images of chol-PK organoids with *Prss22, Lamc2* and *Plau* deletions, respectively. The CRISPR/Cas9-induced genetic modifications in the targeted genes are indicated below. (C) Scatter plot depicting mRNA expression of *Prss22, Lamc2* and *Plau* in chol-PK (n=3) and the derived knockout organoids (n=4). (D) Dot plot measuring proliferation of the indicated organoid lines via bioluminescence at the depicted time points. (E) Liver tumor weight (E), and liver/body weight ratio (F) for both chol-PK and hep-PK derived liver tumors plotted against harvest time point as weeks post implantation. Data points represent individual mice. Statistical significance was defined as: *p* < 0.05 (*), *p* < 0.01 (**), *p* < 0.001 (***), and *p* < 0.0001 (****).

## Supplementary Tables

STable 1. List of 1583 upregulated genes in chol-PK organoids.

STable 2. List of sgRNA oligos and genomic DNA primers.

STable 3. List of antibodies for IHC.

STable 4. List of antibodies for MIBI.

STable 5. List of primer sequences for RT-qPCR.

